# EphB1 controls proper long-range cortical axon guidance through a cell non-autonomous role in GABAergic cells

**DOI:** 10.1101/2022.02.28.482352

**Authors:** Ahlem Assali, George Chenaux, Jennifer Y. Cho, Stefano Berto, Nathan A. Ehrlich, Christopher W. Cowan

**Author notes:** ICM - Brain Institute, Paris, France.

## Abstract

EphB1 is required for proper guidance of cortical axon projections during brain development, but how EphB1 regulates this process remains unclear. We show here that *EphB*1 conditional knockout (cKO) in GABAergic cells (*Vgat*-Cre or *Dlx1/2*-Cre), but not in cortical excitatory neurons (*Emx1*-Cre), reproduced the cortical axon guidance defects observed in global *EphB1* KO mice. Interestingly, in *EphB1* cKO^Vgat^ mice, the misguided axon bundles contained comingled striatal GABAergic and somatosensory cortical glutamatergic axons. In wildtype mice, somatosensory axons also co-fasciculated with striatal axons notably in the globus pallidus, suggesting that a subset of glutamatergic cortical axons normally follows long-range GABAergic axons to reach their targets. Surprisingly, the ectopic axons in *EphB1* KO mice were juxtaposed to major blood vessels. However, conditional loss of *EphB1* in endothelial cells (Tie2-Cre), or in mural and oligodendrocyte precursor cells (Cspg4-Cre) did not produce the axon guidance defects, suggesting that EphB1 in GABAergic neurons normally promotes avoidance of these ectopic axons from following the developing vasculature. Together, our data reveal a new role for EphB1 in GABAergic neurons to influence proper cortical glutamatergic axon guidance during brain development.

## INTRODUCTION

In the developing nervous system, binding of EphB tyrosine kinase receptors to their cell surface-localized ephrin “ligands” triggers bidirectional, intracellular signaling events that regulate proper axon guidance, cell migration, synapse formation, and synapse plasticity [1]. Previous studies revealed deep-layer cortical axon guidance defects in the ventral telencephalon (VTel) of *EphB1*^-/-^ mice [2–4], and these guidance errors were exacerbated in the *EphB1*^-/-^;*EphB2*^-/-^ mice, suggesting partial compensation by EphB2 receptors [2]. EphB1 shows peak expression in cortical layers V and VI at mouse embryonic day 15.5 (E15.5) [2, 4] – a time when long-range cortical axons are navigating toward their subcortical target zones [5]. However, at this developmental stage, EphB1 is also expressed in the developing epithalamus [2], GABAergic cell progenitor structures (i.e., ganglionic eminences and the preoptic area), GABAergic interneurons migrating toward the developing cortex [6], and GABAergic spiny projection neurons (SPNs) of the developing striatum. Developing deep-layer cortical axons also express ephrin-Bs and navigate through EphB1-expressing brain regions in the VTel. As such, it was unclear whether EphB1 regulates cortical axon guidance in a cortical cell autonomous manner or whether EphB1 regulates long-range cortical glutamatergic projections indirectly via a key role in other cell populations.

To investigate the cell populations in which EphB1 regulates proper cortical axon guidance, we generated a new floxed *EphB1* mouse to allow for Cre-dependent *EphB1* loss-of-function analysis. Our findings reveal that EphB1 is required within Vgat (vesicular GABA transporter)-positive cells, but surprisingly not in glutamatergic cortical neurons, to control long-range cortical axon guidance in the VTel. Moreover, in *EphB1* KO mice, we observed numerous aberrant subcortical axon bundles comprising both GABAergic striatal axons and glutamatergic cortical axons, suggesting that a subset of long-range cortical axons normally fasciculate along a subpopulation of long-range GABAergic axons to reach their proper target(s). Surprisingly, the misguided axons preferentially grew along major blood vessels in the VTel, suggesting that axonal EphB1 functions to repel navigating axons away from the developing blood vessels. These effects were not produced by *EphB1* loss-of-function in D1 or D2 dopamine receptor-expressing SPNs, Tie2-expressing vascular endothelial cells, or Cspg4-expressing mural and oligodendrocyte precursor cell populations, suggesting that EphB1 functions in a subpopulation of Vgat-positive neurons to indirectly produce cortical axon guidance defects in apposition to developing striatal vasculature.

## MATERIAL AND METHODS

### Animals

*EphB1* knockout mice (*EphB1*^-/-^) on a C57BL/6 background were generated by crossing *EphB1*^-/-^ mice on a CD-1 background (donated by Dr. Henkemeyer, [7]) to C57BL/6 background strain (>8 generations). *EphB1*-LacZ mice were donated by Dr. Henkemeyer [8]. *EphB1*^ΔEx3/ΔEx3^ mice (*EphB1* total loss-of-function) were generated by crossing floxed *EphB1* mice (*EphB1*^lox/lox^, described below) to Prm-Cre mice (Jackson Laboratory #003328) to produce germline recombination. The Prm-Cre allele was subsequently removed during repeated backcrossing to C57BL/6J wild-type mice. *EphB*1 conditional knockout mice (*EphB1* cKO) were generated by crossing *EphB1*^lox/lox^ mice with cell type-selective Cre-expressing transgenic mice (Emx1-Cre (Jackson Laboratory #005628), Vgat-Cre (Jackson Laboratory #028862), dlx1/2-Cre (I12b-Cre donated by Dr. Rubenstein, Potter et al, 2009), Somatostatin-Cre (Sst-Cre, Jackson Laboratory #013044), Drd1a-Cre (Jackson Laboratory #37156-JAX), Drd2-Cre (GENSAT; MMRRC # 036716-UCD), Tie2-Cre (Jackson Laboratory # 008863), parvalbumin-Cre (PV-Cre, Jackson Laboratory # 017320), Nkx2.1-Cre (Jackson Laboratory #008661), Cspg4-Cre (donated by Dr. Nishiyama), and they were compared to their Cre-negative littermate controls. The Vgat-Cre, Tie2-Cre, PV-Cre, Drd1-Cre and Drd2-Cre lines were crossed to the Ai14 reporter mouse line (Jackson Laboratory #007914). To visualize long-range GABAergic projections, the Vgat-Cre line was crossed to both *EphB1*^lox/lox^ and Ai14 (*EphB1* cKO^Vgat^; tdTomato^Vgat^) mice or to Ai14 (tdTomato^Vgat^). To visualize thalamic and striatal axons in *EphB1* mice, Gbx-CreER^T2^-IRES-EGFP (Jackson Laboratory #022135), Drd1-tdTomato (Jackson Laboratory #016204) and Drd2-GFP (Mouse Genome Informatics MGI:3843608) reporter mice were crossed to *EphB1*^-/-^ mice or to *EphB1*^ΔEx3/ΔEx3^ mice. To visualize the projections from PV-positive neurons, the PV*-*tdTomato reporter line (Jackson Laboratory, #027395) was crossed to *EphB1*^ΔEx3/ΔEx3^ mice. All procedures were conducted in accordance with the MUSC Institutional Animal Care and Use Committee (IACUC) and NIH guidelines.

### Generation of floxed EphB1 mutant mice (EphB^lox/lox^; Suppl. Figure 1)

A targeting vector was generated by using a pL452 based mini targeting vector [9], which was recombined into mouse 129 strain bacterial artificial chromosome clone BMQ422J21 [10]. The mini targeting vector was cloned so that loxP site #1 was inserted 722 bp upstream of EphB1 exon 3 and loxP site #2 is 293 bp downstream of exon 3. In addition to this, an FRT site flanked insertion was cloned downstream of loxP site #1, which contained the following cassettes: an engrailed 2 slice acceptor, the internal ribosomal entry site (IRES) from encephalomyocarditis virus (EMCV) [11], a bovine tau protein & enhanced green fluorescent protein (eGFP) expression sequence [12], an SV40 polyadenylation signal, and a neomycin selection cassette driven by prokaryotic and eukaryotic constitutive promoters. A capture vector as used to retrieve 10,630 bp for the left homology arm and 10,146 bp is for the right homology arm with flanking diphtheria toxin and thymidine kinase negative selection cassettes, respectively. Murine stem cells were targeted and screened by the UC Davis Mouse Biology Program. Mice were crossed with a Flp recombinase germline expression mice to remove the FRT flanked knock-in cassettes to generate *EphB1*^lox/lox^ mice lacking the selection cassettes. Cells that express Cre recombinase delete 1854 bp of the *EphB1* genomic locus, which includes all of exon 3, to generate EphB1^Δexon3^ mice. Exons 1 and 2 have the potential to generate a small, truncated protein. In the *EphB1*^Δexon3^ mice, exon 2 splicing to exon 4 is predicted to produce either nonsense mediated decay or to code for the following protein sequence: MALDCLLLFLLASAVAAMEETLMDTRTATAELGWTANPASG**PVLRGPSRPARKLKAA PTAPPTVAPLQRRLPSAPAGLAITELTLIHQRWRVLVSHRVLEMSSPS,** Bolded letters are produced by a frame-shift and represent the predicted non-EphB1 amino acids prior to the premature stop codon. The primers used to genotype floxed *EphB1* mice were: forward primer 5’GGGAGAAGAGAGAGCCTAC3’, and reverse primer 5’CCAGAGGGCTTTGAGTTAAT3’ (floxed band: 316bp; wild-type band: 420bp).

### Immunohistochemistry

Adult mice were anesthetized with Ketamine/Xylazine diluted in 0.9% saline (120 mg/kg and 16 mg/kg, respectively) and hypothermia anesthesia was used for pups by placing them in ice for 5-8 minutes. Postnatal day 0 (P0) pups and adult mice were perfused transcardially with 4% (w/v) paraformaldehyde (PFA) in phosphate buffered saline (PBS), the brains were post-fixed overnight in 4% PFA, cryoprotected in 30% sucrose, and then sectioned at 40 μm (adults) and 70 μm (pups) using a cryotome. Sections were washed in PBS, incubated in blocking solution (5% (v/v) normal donkey serum, 1% (w/v) Bovine Serum Albumin, 0.2% (v/v) glycine, 0.2% (w/v) lysine, 0.3% (v/v) Triton X-100 in PBS) for 1 hour at room temperature (RT) with shaking, incubated with primary antibodies diluted in blocking solution (rat anti-L1CAM (1:1000) from MilliporeSigma #MAB5272; chicken anti-GFP (1:1000) from Abcam #13970; mouse anti-Brn3a (1:200) from Fisher #MAB1585; rabbit anti-DsRed (for tdTomato staining; (1:1000)) from Living Colors #632496; Rabbit anti-RFP (for tdTomato staining; (1:3000)) from Rockland #600-401-379; goat anti-CD31 (1:800) from Novus Biologicals #AF3628; mouse anti-elastin (1:800) from Sigma #MAB2503) overnight at 4C under shaking, washed 3 times for 10 minutes in PBS with gentle shaking at RT, incubated with secondary antibodies diluted in blocking solution (Cy3 donkey anti-rat (1:500) from Fisher #NC0236073; AlexaFluor 488 donkey anti-mouse (1:500) from Fisher #NC0192065; AlexaFluor 488 donkey anti-chicken (1:500) from Fisher #703-545-155) for 90 minutes at RT with shaking and protected from light, washed 3 times for 10 minutes in PBS, and mounted in ProLong Gold Antifade Mountant (Invitrogen #P36931).

### XGal staining

Adult mice were perfused transcardially with 4%(w/v) PFA, and the brains were post-fixed overnight in 4%(w/v) PFA. Female mice were bred and vaginal plugs were assessed with the day of plug detection considered as embryonic day 0.5 (E0.5). After live-decapitation, embryo brains were drop-fixed in 4% PFA overnight. Adult and embryo brains were then cryoprotected in 30% sucrose. Embryo brains were plunged in M1-embedding matrix (ThermoFisher #1310), flash-frozen for 1 minute in isopentane between -20°C and -30°C, and stored at -80°C. Adult brains were cut at 40μm using a cryotome and embryo brains at 20μm using a cryostat. XGal staining was performed using the beta-galactosidase staining kit (Mirus #MIR 2600), following the provided protocol. Briefly, the sections were washed in PBS, incubated in the Cell Staining Working Solution containing the X-Gal Reagent in a dark, humidified chamber at 37°C overnight, washed once in PBS, and mounted in ProLong Gold Antifade Mountant (Invitrogen #P36931).

### Myelin Stain

Sections were stained for myelin using the BrainStain Imaging kit (ThermoFisher #B34650; FluoroMyelin, 1:300), following the vendor protocol.

### RT-PCR

RNA extraction was performed using the miRNeasy Mini kit (Qiagen #1038703), following the kit’s protocol. Total RNA was reverse-transcribed using Superscript III (Invitrogen) with random hexamers, following the kit’s protocol. PCRs were performed using the complementary DNA to detect *EphB1* expression (for *EphB1* cKO^Emx^: forward primer 5’ TACAGAGATGCGCTT 3’, reverse primer 5’ ACAGCGTGGCCTGCA 3’; for *EphB1^ΔEx3/ΔEx3^* and for *EphB1* cKO^Vgat^: forward primer 5’ AGACATTGATGGACACAAGG 3’, reverse primer 5’ TCAAAGTCAGCTCGGTAATA 3’). GAPDH was used as a control (forward primer 5’ TGAAGGTCGGTGTCAACGGATTTGGC 3’; reverse primer 5’ CATGTAGGCCATGAGGTCCACCAC 3’).

### RNAscope^®^

After live-decapitation, the brains were plunged in M1-embedding matrix (ThermoFisher #1310), flash-frozen for 1 minute in isopentane between -20°C and -30°C, stored at -80°C, and then cut at 16 μm thick slices using a cryostat. Sections were kept at -20°C during all the cutting process and stored at -80°C. RNAscope^®^ was performed using the RNAscope^®^ kit (ACD #323110) and following the ACD protocol provided by the manufacturer. Sections were immersed in 4% (w/v) PFA for 15 minutes, then in 50% (v/v) EtOH for 5 minutes, then in 70% (v/v) EtOH for 5 minutes, and then twice in 100% EtOH for 5 minutes. Sections were covered by RNAscope^®^ Hydrogen Peroxide for 10 minutes in a humidified chamber and washed 3 times in PBS, and the protease incubation step was omitted. Sections were placed in a humidified chamber in the HybEZ^TM^ Oven for all the following steps at 40°C and were washed twice for 2 minutes in Wash Buffer after each of the following incubation steps. Sections were incubated with the probes (ACD EphB1 custom probe designed in EphB1 exon3 #541171-C2; Vgat probe #319191; tdTomato probe #317041-C3; C1 probes were used without dilution; C2 and C3 probes were diluted 50 times in C1 probe or in Diluent) and placed at 40°C for 2 hours. AMP1, AMP2, and AMP3 were successively applied on the sections for 30 minutes each at 40°C. Sections were then covered by the appropriate HRP-C and placed at 40°C for 15 minutes, covered by the appropriate fluorophore (PerkinElmer #NEL744E001KT (Cyanine3); #NEL741E001KT (Fluorescein); #NEL745E001KT (Cyanine5); 1:2000 in TSA Buffer provided in RNAScope^®^ kit), placed at 40°C for 30 minutes, covered by the HRP blocker and incubated at 40°C for 15 minutes. Finally, DAPI was applied for 1 minute and sections were mounted in ProLong Gold Mountant (Invitrogen #P36931).

### Stereotaxic injections

Unilateral stereotaxic injections of AAV5-CaMKIIα-EGFP (Addgene #50469; virus titer ≥ 3×10¹² vg/mL) and AAV5-hSyn-DIO-hM4D(Gi)-mCherry (Addgene #44362; virus titer ≥ 7×10¹² vg/mL) viruses were performed on adult anesthetized (isoflurane) control (Vgat-Cre) and *EphB1* cKO^Vgat^ (*EphB1*^lox/lox^ xVgat-Cre) mice, into the dorsal striatum (DV: -2.8, ML: +1.6, AP: 0; 150 nL) and into two locations of the somatosensory cortex (1. DV: -1.9, ML: +3.2, AP: -0.4; 2. DV: -1.4, ML: +2.7, AP: -1.7; 200 nL in each location), using a Nanoinjector (FisherScientific #13-681-455; 50nL/30sec). Placement was confirmed by immunohistochemistry (GFP and DsRed antibodies; see above).

### Bioinformatics analysis of a single cell RNAseq dataset from whole mouse embryonic brain [13]

Single cell gene expression data was download from http://mousebrain.org/ (PMID: 34321664). Briefly, loom files were converted into Seurat objects with UMI data and meta data. Downstream analysis was performed in R with Seurat (v4.1.0) (PMID: 34062119) and customized R scripts. Data were preprocessed, normalized, and filtered. The Seurat function DotPlot was used to visualize the gene expression and relative abundance of the genes of interest across the different cell subclasses. Cell information was stored in the meta data.

## RESULTS

### Cortical axon guidance defects in *EphB1* KO mice

Since the background strain can influence phenotypes in mutant mice, we backcrossed the *EphB1*^-/-^ mice (on a CD-1 background; [7]) to C57BL/6 background strain (>8 generations). Similar to the *EphB1*^-/-^ on the CD-1 strain [2], the BL/6-backcrossed *EphB1*^-/-^ mice showed multiple axon guidance defects at postnatal day 0 (P0) (Fig. 1), including disorganized axons and ectopic axon bundles within the dorsal striatum (Fig. 1*B* and *C*; see arrows), descending ectopic axon bundles within the internal capsule, and ectopic axon projections descending from the external capsule and terminating near the brain floor (Fig. 1*B* and *D*; see arrows). Previous studies also showed that axons from the somatosensory cortex aberrantly grow in the posterior branch of the anterior commissure in *EphB1*^-/-^ mice [4].

**Figure 1:**
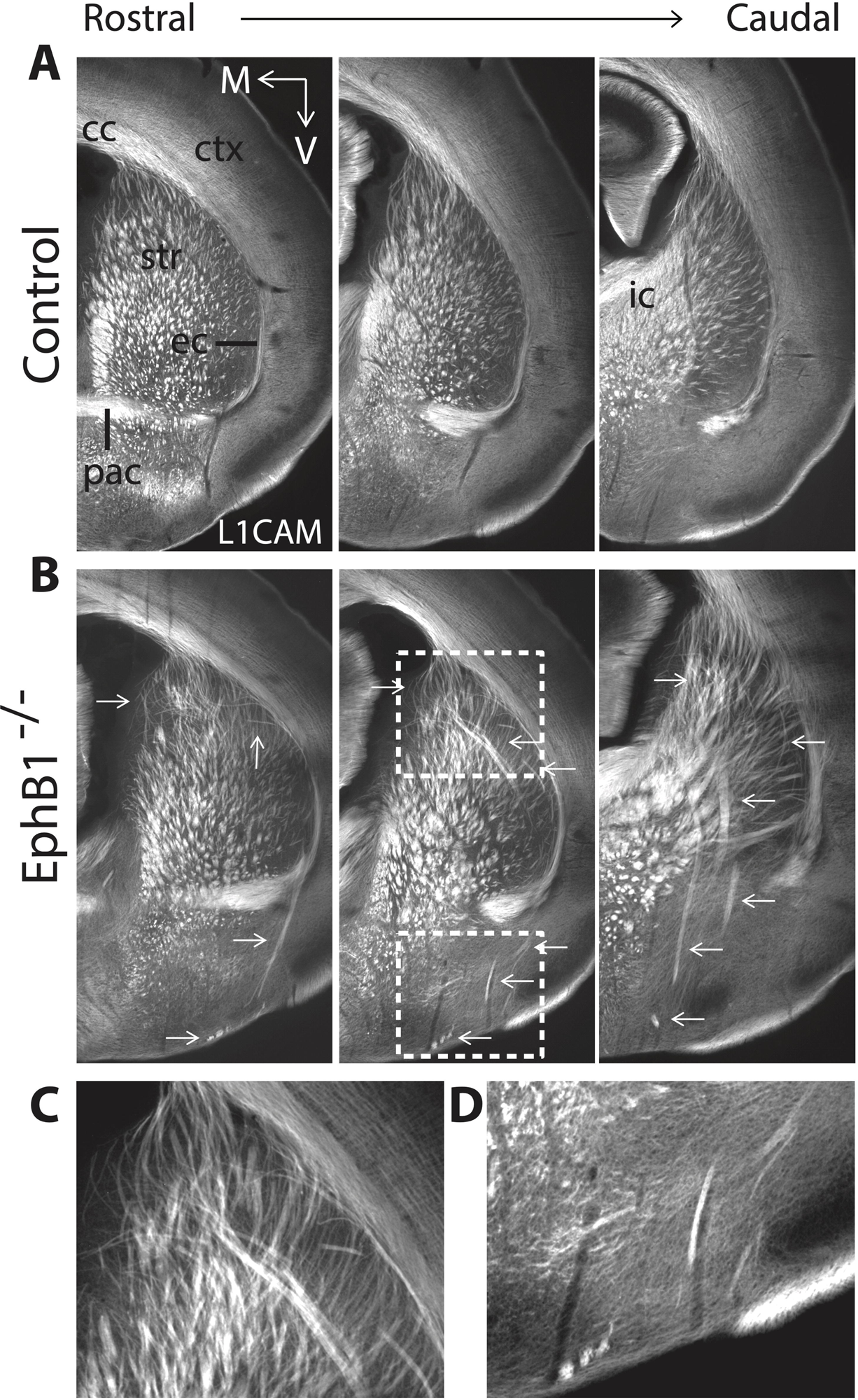
Axon guidance defects in *EphB1*^-/-^ mice on a C57Bl6/J background. **A, B.** L1CAM staining on coronal sections at three different rostro-caudal levels at P0 in control mice (**A**), and in *EphB1*^-/-^ mice (**B**), with abnormal axon bundles in the dorsal striatum joining the external capsule and posterior branch of the anterior commissure, and terminating in the ventral part of the brain, as well as abnormal axon bundles in the internal capsule. **C, D.** Magnified images corresponding to the white squares in **B**. The images were taken using a microscope 10X objective. ctx: cortex; str: striatum; cc: corpus callosum; ec: external capsule; ic: internal capsule; pac: posterior branch of the anterior commissure. V: ventral; M: medial. n = 5 *EphB1*^-/-^ and 8 control littermates.

### Loss of *EphB1* does not influence thalamic axon guidance

We previously reported subtle deficits in thalamic axon guidance in *EphB1*^-/-^;*EphB2*^-/-^ mice [2]. Since misguided VTel axons in *EphB1*^-/-^ mice originate largely from the cortex [2, 4] and ascending thalamocortical projections are thought to influence corticothalamic axon guidance [14], we analyzed *Gbx*-expressing thalamocortical projections in the *EphB1*^-/-^ mice. In the *EphB1*^-/-^ mice, the Gbx-GFP-labeled axon projections appeared normal (Suppl. Fig. 2*A* and *B*), the L1CAM-positive misguided axons did not co-localize with GFP-positive thalamic fibers (Suppl. Fig. 2*C* and *C’*), and the cortical barrel fields were similar to wild-type controls (Suppl. Fig. 2*D* and *E*), suggesting that the cortical axon guidance errors in the VTel of *EphB1*^-/-^ mice are unlikely to be caused by aberrant thalamocortical axon navigation.

EphB1 is highly expressed in a dorsal region of the early developing thalamus [2]. Careful analysis of EphB1 expression (using Xgal staining in EphB1-lacZ mice) in the Gbx-GFP;EphB1-lacZ mice showed that EphB1 is highly expressed in the epithalamus (Suppl. Fig. 3B), but largely undetectable in the developing thalamus (Suppl. Fig. 3*A*). Indeed, the habenula-specific marker Brn3a [15] revealed strong co-localization with EphB1 (X-gal) at E14.5, and EphB1 was highly expressed in the adult habenula (X-gal; Suppl. Fig. 3*B* and *C*, respectively). Despite its strong expression in habenula, we detected no axon guidance defects in the habenular commissure, the fasciculus retroflexus, or the habenular axon tract in the interpeduncular nucleus in P0 *EphB1*^-/-^ pups (Suppl. Fig. 3D). Together these findings suggest that EphB1 is expressed highly in the developing epithalamus/habenula, but it’s not required for normal habenular or thalamic axon guidance. The deficits in thalamic axon guidance in *EphB1*^-/-^;*EphB2*^-/-^ mice [2] might be due to the loss of *EphB2* or to a synergistic effect of the loss of both *EphB1* and *EphB2*.

### Generation of an *EphB1* conditional loss-of-function mutant mouse

To determine the cell population(s) in which *EphB1* functions to regulate proper cortical axon guidance, we generated a mutant mouse with loxP sites flanking *EphB1* exon 3 (*EphB1*^lox/lox^; Suppl. Fig. 4; Suppl. Fig. 1) using traditional homologous recombination. To confirm that Cre-dependent loss of *EphB1* exon 3 reproduced the original *EphB1*^-/-^ phenotype, we generated germline transmission of the Δexon 3 allele (Prm1-Cre). After extensive backcrossing to C57BL/6, we confirmed the loss of *EphB1* expression in the brain of *EphB1*^ΔEx3/ΔEx3^ mice (Suppl. Fig. 4A), and importantly, we observed the same VTel axon guidance defects observed in the original *EphB1*^-/-^ mice (Suppl. Fig. 4B).

### *EphB1* regulates cortical axon guidance via a role in GABAergic neurons

Since EphB1 is expressed in the deep layers of the developing cortex [2, 4], we crossed *EphB1*^lox/lox^ mice with Emx1*-*Cre mice to generate conditional *EphB1* KO (*EphB1* cKO^Emx1^) in most cortical and hippocampal glutamatergic pyramidal neurons starting at ∼E11.5 [16]. Despite the loss of cortical EphB1 expression in the *EphB1* cKO^Emx1^ mice (Suppl. Fig. 5A), we observed no cortical axon guidance defects (Figs. 2A and 2B). EphB1 is also highly expressed in multiple GABAergic neuron populations [2, 6], including those within the developing VTel. To test EphB1 function in GABAergic neurons, we generated an *EphB1* cKO in virtually all GABAergic populations using Vgat-ires*-*Cre mice [17] (Suppl. Figs. 5*B, C,* and *D*). Interestingly, P0 *EphB1* cKO^Vgat^ pups completely phenocopied the cortical axon guidance defects found in the global *EphB1*^-/-^ pups (Figs. 2*C* and *D*). We also observed the cortical axon guidance defects in P0 *EphB1* cKO*^dlx1/2^*pups (Fig. 4*A*) – a distinct GABAergic cell-specific Cre transgenic mouse [18]. Together, these data revealed that EphB1 functions in GABAergic cells to influence cortical axon guidance.

**Figure 2:**
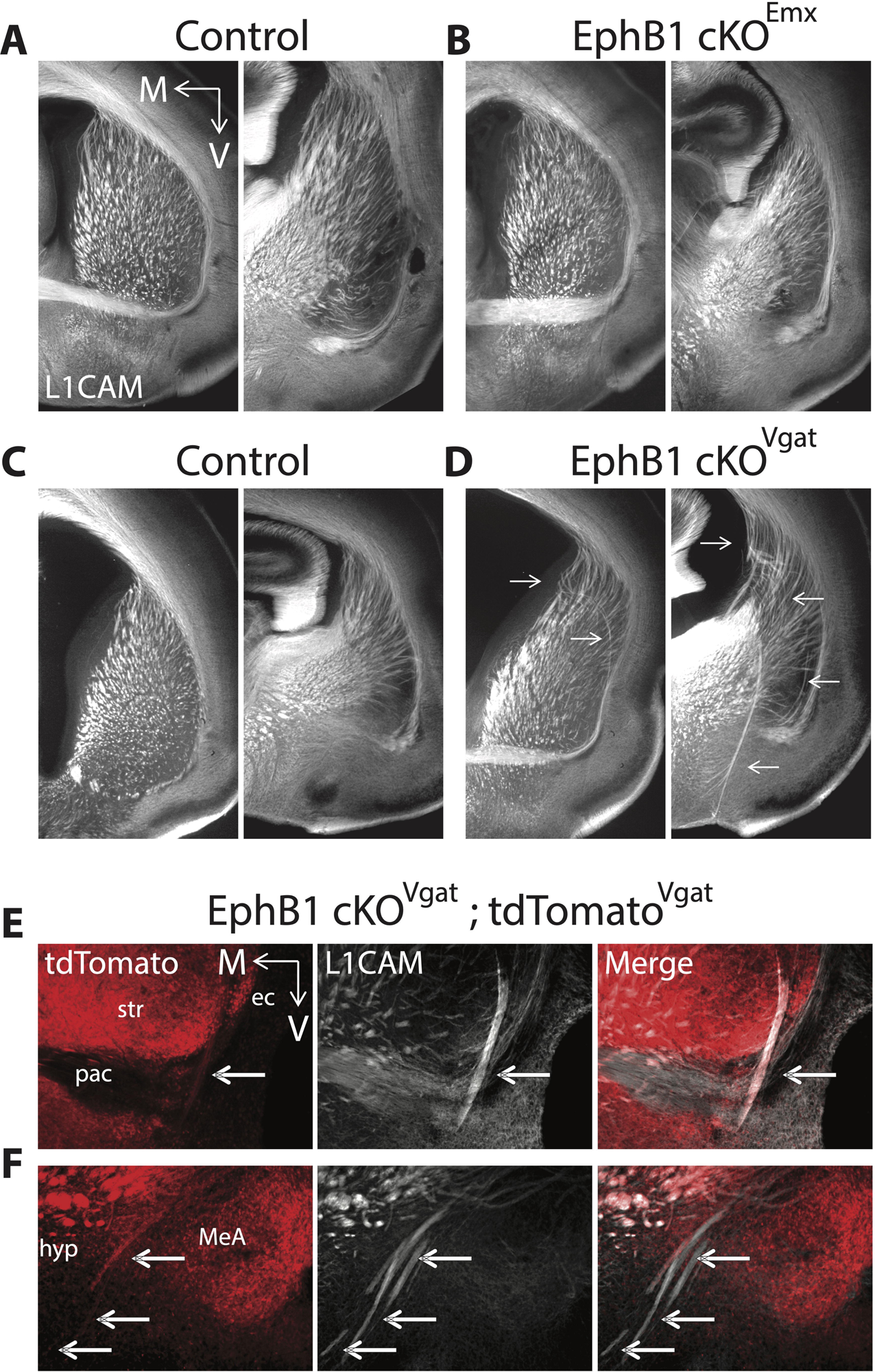
Axon guidance defects after *EphB1* deletion from GABAergic cells. **A-D.** L1CAM staining on coronal sections at two different rostro-caudal levels at P0 in controls (**A, C**), in *EphB1* cKO^Emx^ (**B**), and in *EphB1* cKO^Vgat^ (**D**). *EphB1* deletion from GABAergic cells, but not glutamatergic cells, phenocopies the axon guidance defects (arrows) found in the global *EphB1*^-/-^ mice. **E, F.** L1CAM and tdTomato co-staining on coronal sections at P0 in *EphB1* cKO^Vgat^;tdTomato^Vgat^ mice (Vgat-Cre x *EphB1*^lox/lox^ x Ai14), showing that the misguided axon bundles include long-range GABAergic projections (arrows). The images were taken using a microscope 10X objective. V: ventral; M: medial; ec: external capsule; pac: posterior branch of the anterior commissure; str: striatum; hyp: hypothalamus; MeA: medial amygdala nucleus. n = 4 *EphB1* cKO^Emx^ and 3 control littermates. n = 5 *EphB1* cKO^Vgat^ and 5 control littermates. n = 3 *EphB1* cKO^Vgat^;tdTomato^Vgat^.

### Cortical axons co-fasciculate with misguided striatal GABAergic axons in absence of ***EphB1***

A subset of cortical GABAergic neurons (e.g., some parvalbumin (PV)- or somatostatin (Sst)-positive neurons), project long-range to subcortical regions [19], suggesting a role of EphB1 in long-range projecting cortical GABAergic axon guidance. Using compound mutant mice *EphB1* cKO^Vgat^;tdTomato^Vgat^ (*EphB1*^lox/lox^ x Vgat-Cre x Ai14), we indeed observed tdTomato-positive GABAergic long-range projecting axons located within the L1CAM-positive ectopic axon bundles at both P0 (Figs. 2*E* and *F*) and E15.5 (Suppl. Fig. 6), suggesting that the GABAergic axon guidance deficits emerge as early as the misguided cortical axons. However, using a Cre-dependent virus approach in *EphB1* cKO*^Vgat^* mice, we did not detect Vgat-positive axons from the somatosensory cortex (i.e., the origin of many of the misprojected VTel axons in *EphB1*^-/-^; *EphB2*^-/-^ mice [2]) within the ectopic axon bundles of *EphB1* cKO^Vgat^ mice (Suppl. Fig. 7). These data suggest that the misguided GABAergic axons do not originate from long-range projecting cortical GABAergic neurons in the somatosensory cortex. GABAergic striatal SPNs in the striatum also project long-range, and EphB1 is expressed in GABAergic cells of the developing and adult striatum ([6]; Suppl. Fig. 5D). Using the same Cre-dependent virus approach in *EphB1* cKO*^Vgat^*mice, we clearly detected bundles of ectopic Vgat-positive cell axons originating from the dorsal striatum in the *EphB1* cKO^Vgat^ mice (Fig. 3D), as compared to the control mice (Fig. 3A). To verify that misguided VTel axons in *EphB1* cKO*^Vgat^* mice also originate from cortical glutamatergic neurons, we used a virus labeling approach to label CaMKIIα-positive glutamatergic neurons in the SSCtx. Indeed, in *EphB1* cKO*^Vgat^* mice, we observed misguided, CamKIIα-positive cortical glutamatergic axons in ectopic VTel axon bundles (Fig. 3E), compared to control mice (Fig. 3B). Importantly, in *EphB1* cKO*^Vgat^* mice, we detected co-fasciculation of ectopic Vgat-positive dorsal striatum axons and SSCtx glutamatergic cell axons within the VTel ectopic axon bundles and in the posterior branch of the anterior commissure (Fig. 3F), as compared to control mice (Fig. 3C). Together, these data revealed that EphB1 controls proper long-range glutamatergic cortical axon guidance through a cell non-autonomous role in GABAergic cells. Interestingly, in control mice, we also observed cortical glutamatergic axons co-mingled with striatal GABAergic axons along the rostro-caudal axis, specifically in the tracts projecting to the globus pallidus (Suppl. Fig. 8A-D) and to the substantia nigra (Suppl. Fig. 8E-H) – two major targets of SPNs. These data strongly suggest that cortical glutamatergic axons traveling through the striatum fasciculate with striatal GABAergic axons to reach their proper targets.

**Figure 3:**
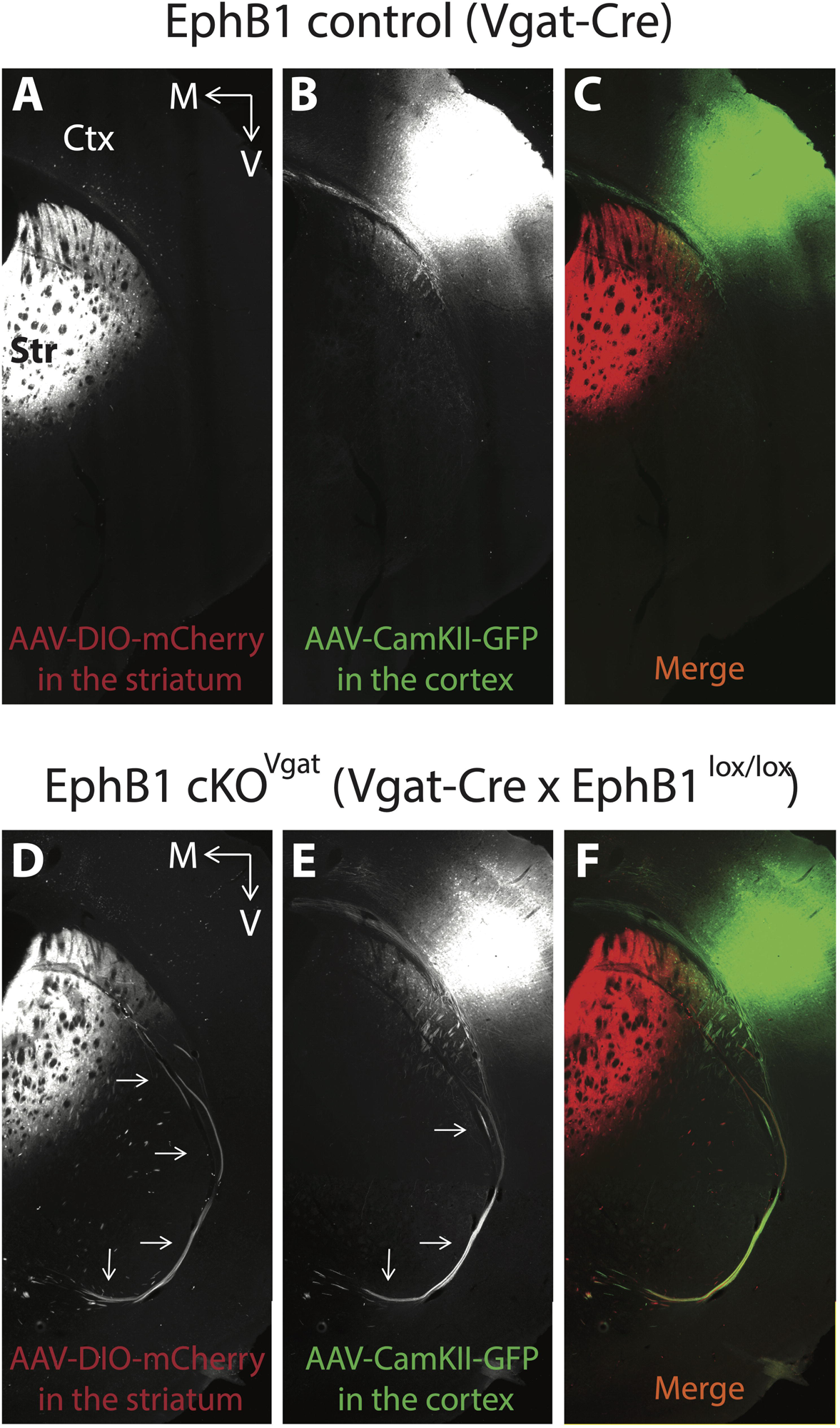
Ectopic cortical glutamatergic projections intermingle with misguided striatal long-range GABAergic projections in *EphB1* cKO^Vgat^ mice. **A-F.** Ds-Red and GFP co-staining on coronal sections in adulthood in control mice (**A-C**) and *EphB1* cKO^Vgat^ mice (**D-F**), following Cre-dependent (DIO) mCherry AAV virus injections in the dorsal striatum (**A**, **D**) and CaMKII GFP AAV virus injections in the somatosensory cortex (**B**, **E**). **C, F.** Merge. The arrows indicate the ectopic axons. The images were taken using a microscope 10X objective. V: ventral; M: medial; Ctx: cortex; Str: striatum. n = 3 *EphB1* cKO^Vgat^ and 3 control littermates.

### Loss of EphB1 in GABAergic D1 or D2 dopamine receptor-expressing populations does not phenocopy the axon guidance deficits observed in *EphB1* KO mice

The vast majority of striatal GABAergic cells are SPNs that express D1 or D2 dopamine receptors [20]. To test whether EphB1 might play a critical role in D1- or D2-SPNs to produce the axon guidance defects, we first analyzed the effect of *EphB1*^-/-^ on D1- and D2-SPN projections in the Drd1-tdTomato or Drd2-GFP reporter mice. However, we failed to detect tdTomato-positive (D1) or GFP-positive (D2) axons within the VTel axon bundles in *EphB1*^-/-^ mice (Suppl. Fig. 9*A* and *B*). Moreover, despite effective recombination in the Drd1-Cre and Drd2-Cre mice at E14.5 (Suppl. Fig. 10A and B), conditional *EphB1* KO in either Drd1-Cre or Drd2-Cre transgenic lines (*EphB1* cKO^D1^ and *EphB1* cKO^D2^) failed to produce the axon guidance deficits (Figs. 4*C* and *D*), suggesting that the VTel axon guidance phenotype in *EphB1*^-/-^ mice is not caused by a key role in developing D1- or D2-SPNs.

**Figure 4:**
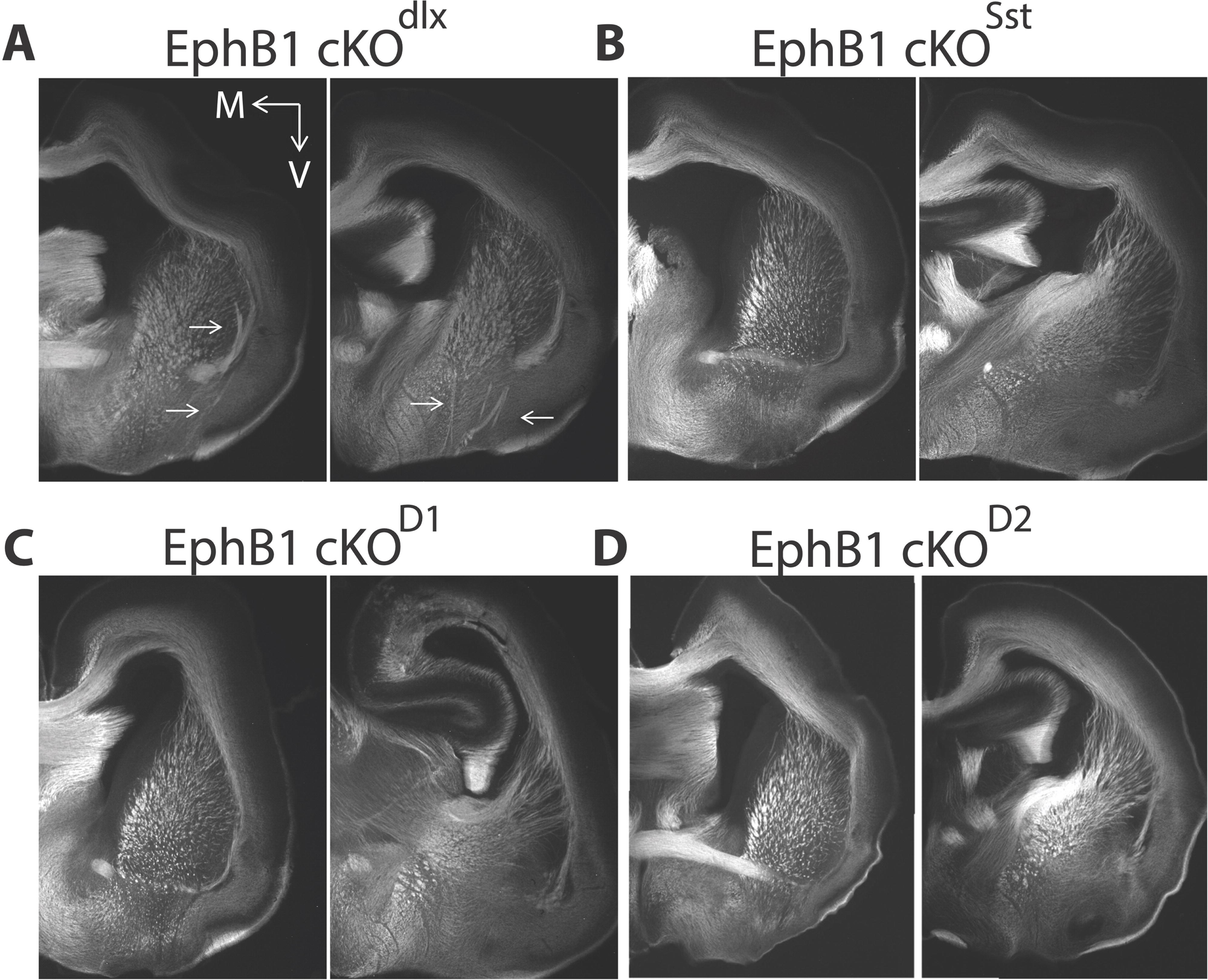
No axon guidance defects after *EphB1* deletion from D1- or D2-SPNs. **A-D.** L1CAM staining on coronal sections at two different rostro-caudal levels at P0 in *EphB1* cKO^dlx1/2^ (**A**), in *EphB1* cKO^Sst^ (**B**), in *EphB1* cKO^D1^ (**C**), and in *EphB1* cKO^D2^ (**D**). The images were taken using a microscope 10X objective. V: ventral; M: medial. n = 4 *EphB1* cKO^dlx1/2^ and 4 control littermates. n = 3 *EphB1* cKO^Sst^ and 3 control littermates. n = 3 *EphB1* cKO^D1^ and 3 control littermates. n = 3 *EphB1* cKO^D2^ and 3 control littermates.

A sub-population of Sst and PV GABAergic neurons are known to project long-range from the cortex ([19] and Fig. 5*A*). PV neurons also project long-range from the globus pallidus and substantia nigra ([21] [22]). In *EphB1^Δ^*^Ex3/ΔEx3^;PV-tdTomato mice, we detected long-range PV-positive axons within the misguided axon bundles (Fig. 5*B*). To test whether *EphB1* plays a key role in PV neurons to produce the cortical and striatal axon guidance errors in the *EphB1* cKO^Vgat^ mice, we used the Nkx2.1-Cre mouse line, that recombines in both PV- and Sst-positive neurons around E10.5 [23]. However, we observed no axon guidance defects in either *EphB1* cKO^Sst^ mice or *EphB1* cKO^Nkx2.1^ mice (Fig. 4B and 5C).

**Figure 5:**
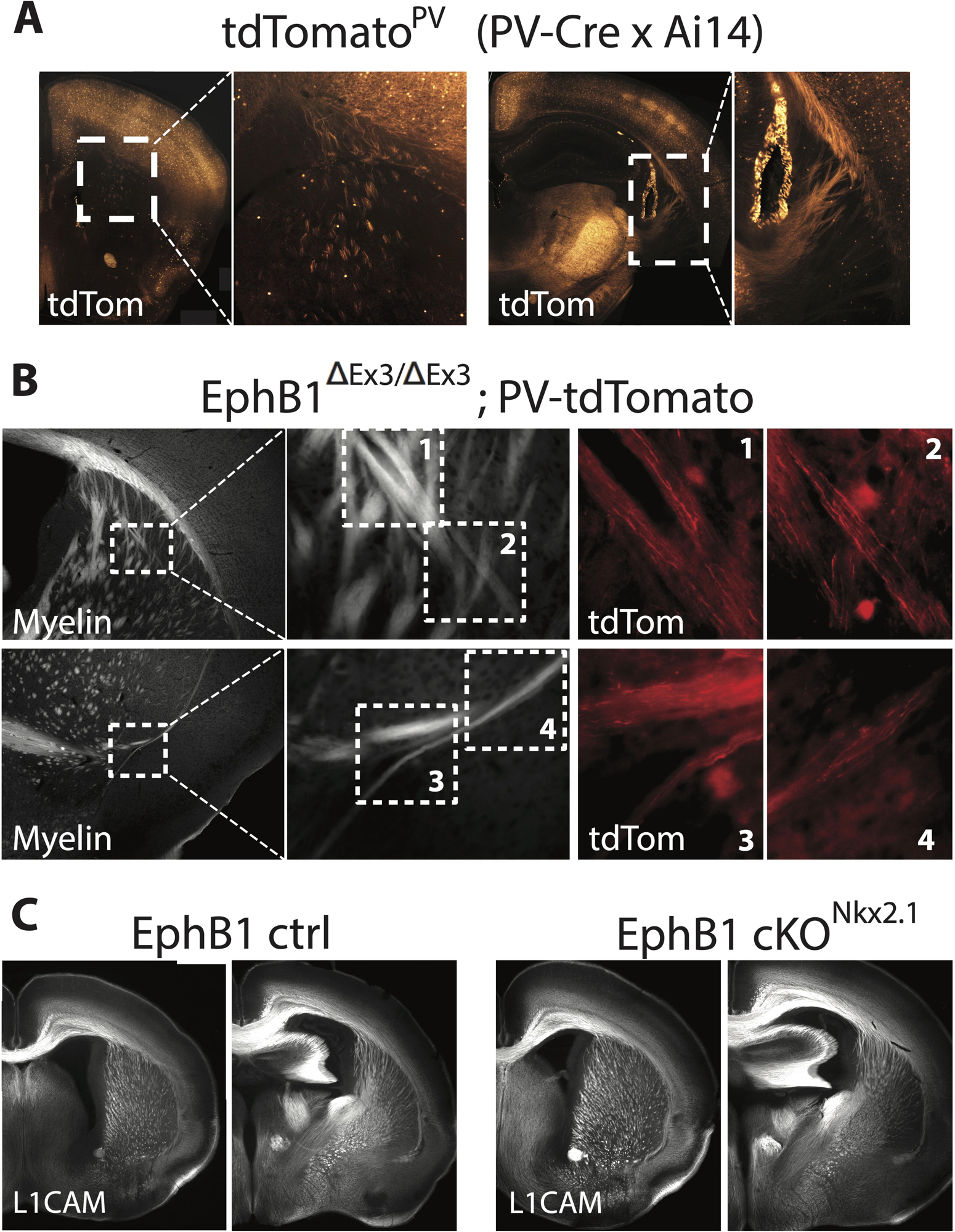
Misguided axons in EphB1-lacking mice include long-range projections from parvalbumin-positive neurons. **A.** Ds-Red staining on coronal sections at two different rostro-caudal levels in adult tdTomato^PV^ mice (PV-Cre x Ai14), showing long-range projections from parvalbumin-positive neurons, including in the dorsal striatum. **B.** Myelin and Ds-Red co-staining on different coronal sections in adult *EphB1*^ΔEx3/ΔEx3^;PV-tdTomato, showing that the misguided axon bundles include long-range projections from parvalbumin-positive neurons. **C.** L1CAM staining on coronal sections at two different rostro-caudal levels in P0 *EphB1* cKO^Nkx2.1^ pups. The images were taken using a microscope 10X objective. n = 3 *EphB1*^ΔEx3/ΔEx3^;PV-tdTomato. n = 4 *EphB1* cKO^Nkx2.1^ and 2 control littermates.

### Loss of EphB1 produces aberrant axon tracts along blood vessels in the ventral telencephalon

In the *EphB1*^ΔEx3/ΔEx3^ mice, we noticed that the VTel ectoptic axon fascicles were typically located in a similar anatomical location and pattern as VTel vasculature (Fig. 6A and B) [24–26]. Interestingly, the ectopic axon bundles in *EphB1*^ΔEx3/ΔEx3^ mice were observed in close apposition to large CD31-positive blood vessels (adults; Fig. 6C) and elastin-positive arteries (P0; Fig. 6D). In addition, long-range GABAergic projections were observed in the ectopic axon bundles located beside the striatal blood vessels (Fig. 7*D*).

**Figure 6:**
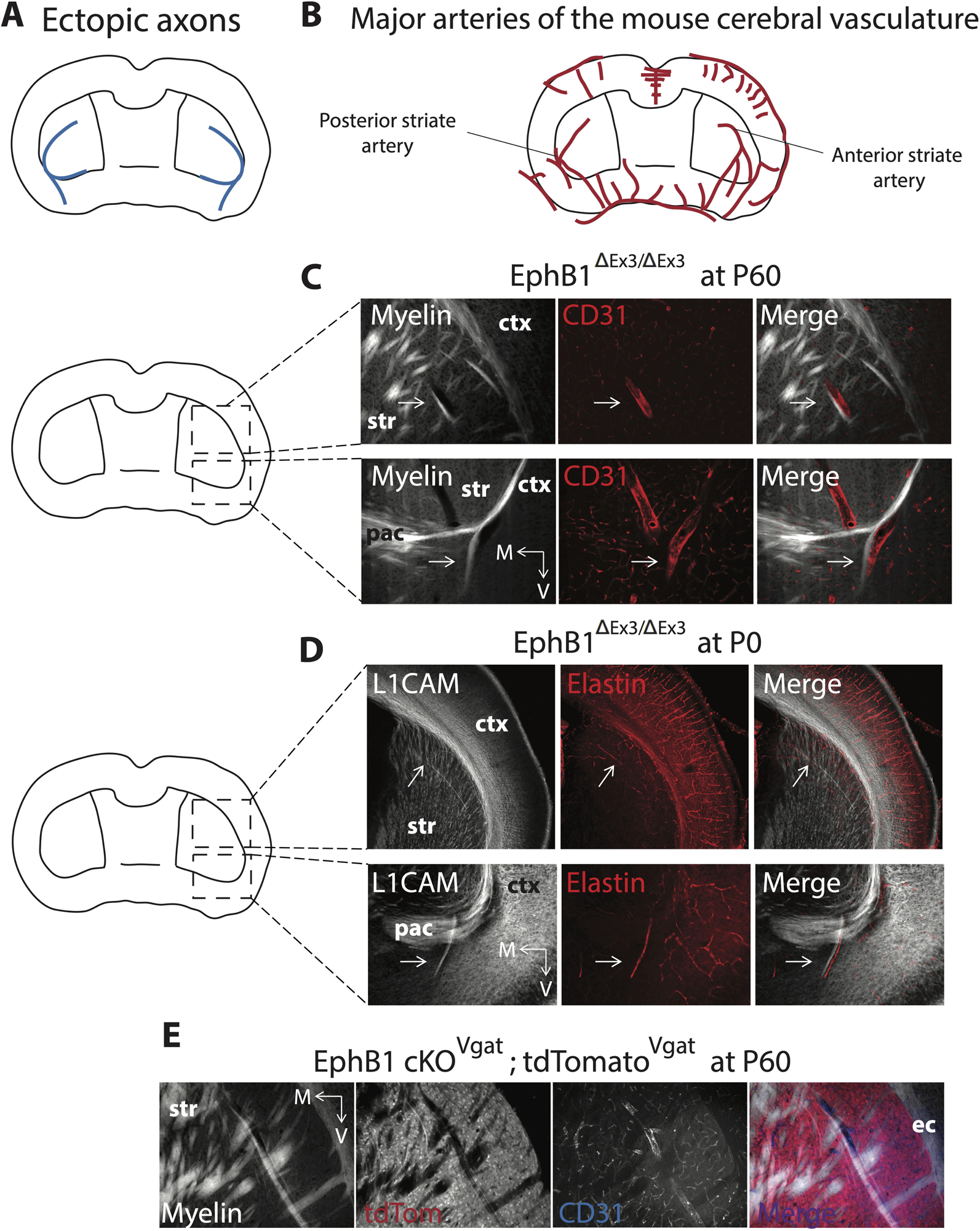
Misguided axons juxtapose with major blood vessels in EphB1-lacking mice. **A.** Scheme representing the ectopic axons after EphB1 deletion. **B.** Scheme representing major arteries of the mouse cerebral vasculature adapted from Dorr et al [24]. **C.** CD31 (endothelial cells marker) and myelin co-staining on different coronal sections in adult *EphB1*^ΔEx3/ΔEx3^ mice. **D.** Elastin (arteries marker) and L1CAM co-staining on different coronal sections in P0 *EphB1*^ΔEx3/ΔEx3^ pups. **E.** CD31, tdTomato, and myelin co-staining on adult *EphB1* cKO^Vgat^;tdTomato^Vgat^ mice (Vgat-Cre x *EphB1*^lox/lox^ x Ai14). V: ventral; M: medial. ctx: cortex; str: striatum; ec: external capsule; pac: posterior branch of the anterior commissure. The images were taken using a microscope 10X objective. n = 3 *EphB1*^ΔEx3/ΔEx3^ at P60 (CD31). n = 3 *EphB1*^ΔEx3/ΔEx3^ at P0 (elastin). n = 3 *EphB1* cKO^Vgat^;tdTomato^Vgat^ at P60.

**Figure 7:**
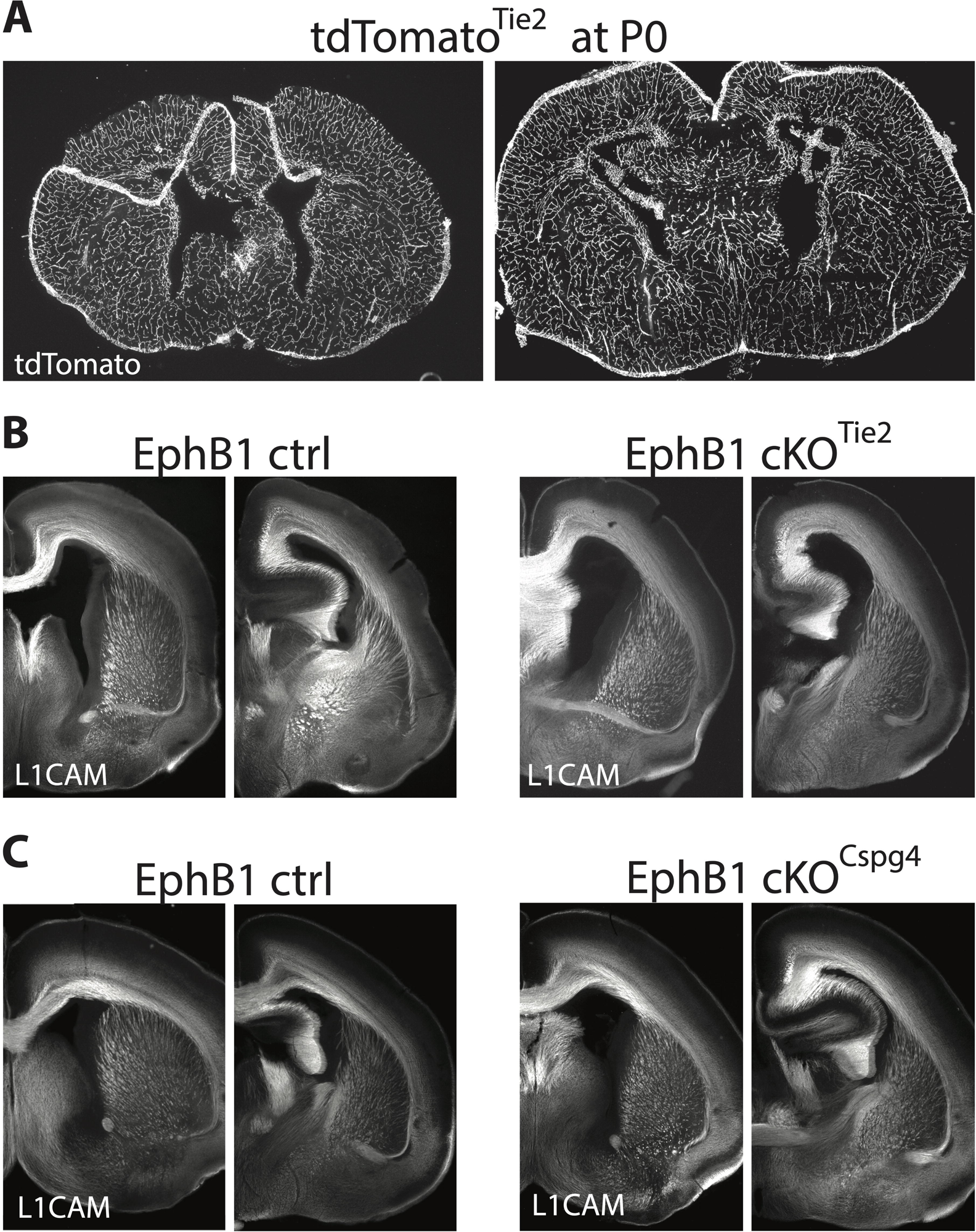
No axon guidance defects after *EphB1* deletion from endothelial, mural and oligodendrocyte precursor cells. **A.** Ds-Red staining on coronal sections at two different rostro-caudal levels in P0 tdTomato^Tie2^ pups (Tie2-Cre x Ai14). **B, C.** L1CAM staining on coronal sections at two different rostro-caudal levels in P0 *EphB1* cKO^Tie2^ pups (**B**, endothelial), and in P0 *EphB1* cKO^Cspg4^ pups (**C**, mural cells and OPCs). The images were taken using a microscope 10X objective. n = 3 *EphB1* cKO^Tie2^ and 3 control littermates. n = 4 *EphB1* cKO^Cspg4^ and 4 control littermates.

Vascular endothelial cells express many axon guidance molecules, including members of the Eph/ephrin family [27, 28] (Suppl. Fig. 11; [13]), that are required for proper vasculature development. Moreover, a subpopulation of endothelial cells also express Vgat and Dlx [29, 30], and EphB1 is expressed in angioblasts (i.e., endothelial cells precursor cells) at E14.5 (Suppl. Fig. 11). To test whether EphB1 is required in endothelial cells to possibly prevent ephrin-B-expressing GABAergic and glutamatergic axons (Suppl. Fig. 11) from following developing blood vessels, we generated vascular endothelial cell-specific *EphB1* cKO ^Tie2^ mice using Tie2-Cre mice [31], where Cre expression in endothelial cells begins by ∼E13 [30]. However, while the Tie2-Cre mice showed specific brain vasculature recombination (Fig. 7A), the *EphB1* cKO^Tie2^ mice displayed no VTel axon guidance phenotypes (Fig. 7B).

Mural cells (vascular smooth muscle cells and pericytes) are found around arteries and veins or microvessels. Oligodendrocyte precursor cells (OPCs) migrate along the vasculature in the developing brain [32]. Both mural cells and OPCs are found associated to endothelial cells as early as E12.5 [33], and NG2 glycoprotein, specifically expressed in OPCs and pericytes, is already present at E13 in the developing mouse brain [34]. In addition, a pool of progenitor cells located in the MGE producing both GABAergic neurons and OPCs, expresses dlx2 as early as E12.5 [35]. According to RNAseq datasets, EphB1 mRNA is expressed in both OPCs and oligodendrocytes [36–38] (Suppl. Fig. 11), suggesting that the phenotype observed in *EphB1* cKO^dlx1/2^ and in *EphB1* cKO^Vgat^ might be due to a loss of *EphB1* from OPCs or mural cells. To determine if the *EphB1*^ΔEx3/ΔEx3^ mice VTel axon guidance defects are produced by *EphB1* deletion in OPCs or mural cells to prevent ephrin-B-expressing GABAergic and glutamatergic axons from following developing blood vessels, we specifically deleted *EphB1* from Cspg4 (NG2)-positive cells. However, the *EphB1* cKO^Cspg4^ mice did not present the axon guidance defects observed in *EphB1*^-/-^ mice (Fig. 7C). Together, these data suggest that EphB1 does not cause the cortical axon guidance deficits via a role in endothelial cells, mural cells, or OPCs, but rather, EphB1 in GABAergic axons promotes avoidance of striatal, and indirectly cortical, axons from the developing brain vasculature (Suppl. Fig. 12 and 13).

## DISCUSSION

To identify the cell populations in which EphB1 is required to regulate cortical long-range axon guidance, we generated and validated a new floxed *EphB1* mouse. This novel tool allowed us to show that, despite its expression in long-range glutamatergic cortical neurons, EphB1 functions in GABAergic cell populations, but surprisingly not in D1 or D2 receptor-expressing striatal SPNs, to influence the proper cortical glutamatergic axon guidance in the developing striatum. We also detected striatal GABAergic axons co-fasciculated with cortical glutamatergic ectopic axon bundles, indicating that EphB1 is required for striatal cell long-range axon guidance, and suggesting that cortical glutamatergic axons are misrouted as an indirect consequence. The Eph/ephrin system is known to be involved in the proper migration of GABAergic neurons generated in the ganglionic eminences and preoptic area [6, 39–43]; however, little was known about the role of the Eph/ephrin system in GABAergic axon guidance. Moreover, the absence of EphB1 in GABAergic cells causes the misguided VTel axons to navigate along the developing striatal vasculature, possibly due to a failure of VTel axons to repel from ephrin-expressing vascular endothelial cells.

Based on the corticothalamic handshake hypothesis [14], and our previous findings in *EphB1*^-/-^;*EphB2*^-/-^ mice showing deficits in both cortical and thalamic axon guidance [2], we speculated that the cortical axon guidance errors were an indirect effect of a role for EphB1 and EphB2 in ascending thalamocortical axons in the developing VTel. In the current study, we focused on the VTel axon guidance phenotypes in the single *EphB1*^-/-^ mouse, which were less severe than the *EphB1*^-/-^;*EphB2*^-/-^ mice [2]. Using the Gbx-EGFP reporter mouse crossed to *EphB1*^-/-^ mice, we observed VTel axon guidance errors, but normal thalamocortical axon projections and no comingling of L1CAM+ ectopic cortical axons with the GFP-positive thalamic axons. Careful examination revealed that EphB1 is highly expressed in the epithalamus (future habenula), but not detectably in the developing thalamus. However, no habenular axon guidance defects were observed in the *EphB1*^-/-^ mice. These findings suggest that the VTel axon guidance defects in *EphB1*^-/-^ mice are not likely to be caused by defects in thalamocortical axon guidance. However, EphB2 is expressed in the thalamus [2], so future studies in compound mutant mice will be required to investigate how EphB1 and EphB2 cooperate to influence proper thalamocortical axon guidance.

EphB1 mediates many of its biological functions through bidirectional signaling induced by contact-mediated binding to ephrins. EphrinB2 is strongly expressed in the developing cortex and striatum [2], and we previously showed that conditional deletion of *ephrinB2* in Nestin-Cre mice, where recombination includes cortical glutamatergic neurons, did not phenocopy the axon guidance deficits found in *EphB1*^-/-^ mice [2]. Conditional deletion of *ephrinB2* using the FoxG1-Cre line produced thalamo-cortical axon guidance deficits similar to those observed in *EphB1*^-/-^;*EphB2*^-/-^ mice, but did not phenocopy the cortical axon guidance deficits found in *EphB1*^-/-^ mice and in *EphB1*^-/-^;*EphB2*^-/-^ mice. EphrinB3, also expressed in the cortex and hippocampus, has been shown to control corticospinal tract axon guidance [44], but future studies will be needed to assess its possible role in the *EphB1* cKO^Vgat^ phenotype.

Using viral-mediated axon tracing tools, we detected GABAergic axons intermingled with the misguided cortical axon bundles in the VTel and located along large blood vessels. Since EphB1 is not required within excitatory cortical neurons for the axon guidance phenotypes but is necessary in Vgat and Dlx1/2-expressing cells, our findings suggest that EphB1 functions in a subset of GABAergic neurons to control proper long-range GABAergic axon guidance and avoidance of the developing VTel vasculature. Moreover, the cortical glutamatergic axon guidance defect is likely caused by co-fasciculation of the descending cortical glutamatergic neuron axons along the misrouted GABAergic fibers. As such, this suggested that a subset of long-range cortical axons might normally fasciculate along a subpopulation of long-range GABAergic axons to reach their proper target. Indeed, we found in control mice, that somatosensory cortical glutamatergic axons co-fasciculated with striatal GABAergic axons along the rostro-caudal axis, specifically in the axon tracts projecting to the globus pallidus and substantia nigra. A very recent study also showed that corticofugal axons fasciculate with striatal axons [45]. In addition, a subpopulation of axons from the somatosensory and motor cortex projects to the globus pallidus [46], which is the predominant target of striatal SPNs. Moreover, the *EphB1*^-/-^ mice display a reduction in corticospinal tract axons [4], suggesting that the misguided axons are largely corticofugal axons. In control mice, there is a subset of corticofugal axons that project to both the spinal cord and to striatal neurons via axon collaterals [47]. All together, these observations strongly suggest that, in our model, a subset of cortical axons are misguided by following misrouted EphB1-null GABAergic striatal axons.

While D1- and D2-SPNs are the predominant, long-range projecting GABAergic cell type in the striatum, conditional loss of *EphB1* in D1 or D2 dopamine receptor-expressing cells failed to produce axon guidance defects, and no D1 or D2-positive axons were clearly observed in the ectopic bundles. However, a very small immature population of SPNs [20] might be the population giving rise to the misrouted striatal axons labeled by the viral approach. Unpublished single nuclei RNAseq data in the mature striatum from our laboratory revealed small subpopulations of Grm8-positive SPNs that express very low or undetectable levels of D1 or D2 dopamine receptor mRNA (data not shown), so it is possible that these unconventional SPNs utilize EphB1 to control their axon guidance and give rise to the EphB1 loss-of-function phenotypes. Sst-positive interneurons in the striatum can project over long distances within the striatum [48], but conditional *EphB1* cKO^Sst^ failed to produce the *EphB1*^-/-^ phenotypes. In normally developing striatum, the PV-expressing GABAergic interneurons project at short-ranges [48], but it remains possible that loss of EphB1 causes striatal PV-positive cells to misproject. In addition, a sub-population of PV-positive neurons (∼17%) in the globus pallidus projects to the striatum [21], and loss of EphB1 in this population could misroute their axons within the striatum. However, we did not observe axon guidance deficits in EphB1^Nkx2.1^ mice, suggesting that PV cells are not the cause of the axon guidance phenotypes. Finally, since conditional deletion of *EphB1* in either D1-SPNs, D2-SPNs, Sst, or PV neurons alone does not produce axon guidance phenotypes, future studies will be needed to identify the specific GABAergic cell subtype(s) responsible for the *EphB1* KO^Vgat^ axon guidance phenotypes.

We found that a subset of striatal GABAergic and cortical glutamatergic axons in *EphB1*^ΔEx3/ΔEx3^ and *EphB1* cKO^Vgat^ are in close apposition to large blood vessels in the Vtel. However, conditional loss of *EphB1* in Tie2-expresssing vascular endothelial cells or in Cspg4-expressing oligodendrocyte precursor cells (OPCs), mural cells, or pericytes did not produce the noted axon guidance deficits, suggesting a role of EphB1 in GABAergic neurons. The developing vasculature and navigating axons express many of the same guidance molecules, including ephrins and Ephs [27, 49], suggesting that both systems can influence each other. However, the interplay between axon navigation and the developing vasculature has only begun to be examined in the central nervous system [50–53]. Several studies showed that some peripheral nerves (*e.g.* autonomic sympathetic axons) navigate along the vasculature via attractive cues expressed on, or secreted by, blood vessels [54]. Conversely, certain sensory nerves can provide a template to guide arterial patterning [49]. Here, we add to this literature by showing that blood vessels also appear to influence striatal and cortical axon guidance. Except in the external capsule, cortical and striatal axons are generally not found in close apposition to the major blood vessels in the VTel, suggesting that EphB1 likely mediates the repulsion of these VTel axons from developing blood vessels. Using a single cell RNAseq dataset from whole embryonic brains at E14.5, we found that ephrinBs are expressed in numerous brain regions and cell types, including neurons but also angioblasts, endothelial cells, mural cells and perivascular fibroblast-like cells. The arteriole endothelium strongly expresses ephrinB2 [28, 55, 56], which could promote EphB1-dependent repulsion of navigating axons in the VTel. Unfortunately, loss of *ephrinB2* from endothelial cells leads to early embryonic lethality (by E11.5) due to angiogenic defects [28], precluding our ability to examine whether loss of vascular e*phrinB2* produces the VTel axon guidance phenotypes seen in the *EphB1*^-/-^ mice. Moreover, we can’t exclude the possibility that in the *EphB1* mutant mice, the colocalization of axons along the VTel blood vessels might be unrelated to EphB1’s ability to mediate repulsive axon guidance.

Together, our findings reveal that EphB1 controls proper long-range cortical axon guidance through a cell non-autonomous role in GABAergic cells. EphB1 is also required to prevent cortical and striatal axons from extending along developing ventral telencephalic vasculature.

## Acknowledgments

The authors would like to thank Dr. Mark Henkemeyer for sharing the EphB1-lacZ knock-in mice. We also thank Ben Zirlin, Brandon Hughes, Rachel Penrod-Martin, Nicolas Narboux-Neme and Nicolas Renier for invaluable assistance with the project. These studies were supported by NIH MH111464 (to C.W.C.), a SFARI research grant #240332 from the Simons Foundation (to C.W.C.), and the MUSC Proteogenomics Facility (NIH GM103499).

## Author contributions

GC designed, produced, and validated at the genomic level the floxed *EphB1* exon 3 targeting vector. SB performed bioinformatics analysis. JYC performed timed breedings. NAE assisted with the RNAscope experiments. AA performed and analyzed all of the other experiments in this paper. AA and CWC designed experiments, interpreted the data, and wrote the paper.

