## Supplemental Figures for "EphB1 controls proper long-range cortical axon guidance through a cell non-autonomous role in GABAergic cells"

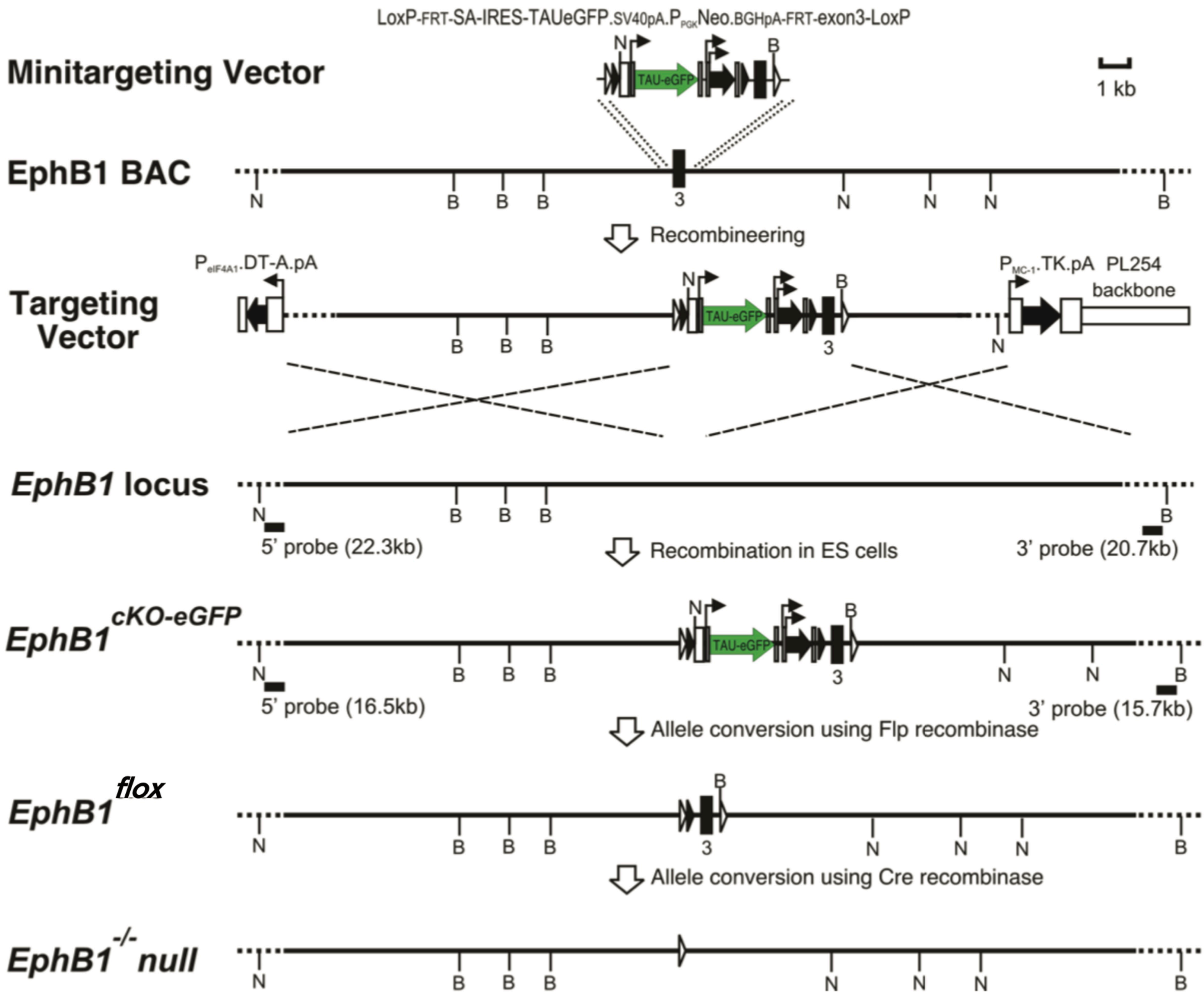


**Supplementary Figure 1**: **Generation of floxed EphB1 mice**. All details are in the materials and methods section.

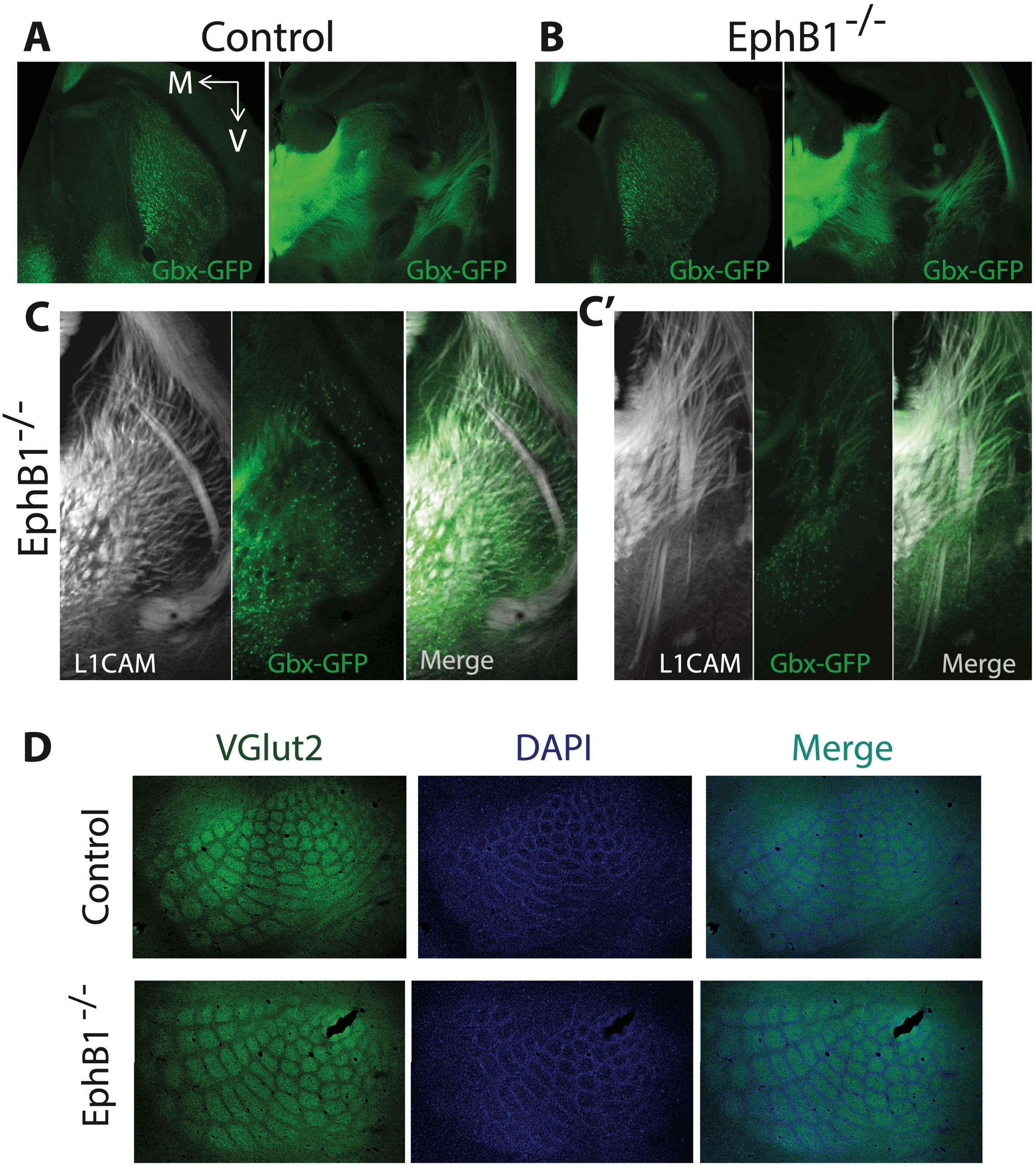


**Supplementary Figure 2**: **No misguided thalamic projections in EphB1$^{-/-}$ mice. A, B.** GFP staining of thalamic nuclei and projections on coronal sections at two different rostro-caudal levels at P0 in Gbx-GFP control (**A**) and in Gbx-GFP; EphB1$^{-/-}$ pups (**B**). **C, C'**. L1CAM and GFP co-staining on coronal sections at two different rostro-caudal levels in P0 Gbx-GFP; EphB1$^{-/-}$ pups. **D, E.** Vglut2 and DAPI co-staining on tangential sections of flattened barrel cortex in adult control (**D**) and EphB1$^{-/-}$ mice (**E**). The images were taken using a microscope 10X objective. V: ventral; M: medial.

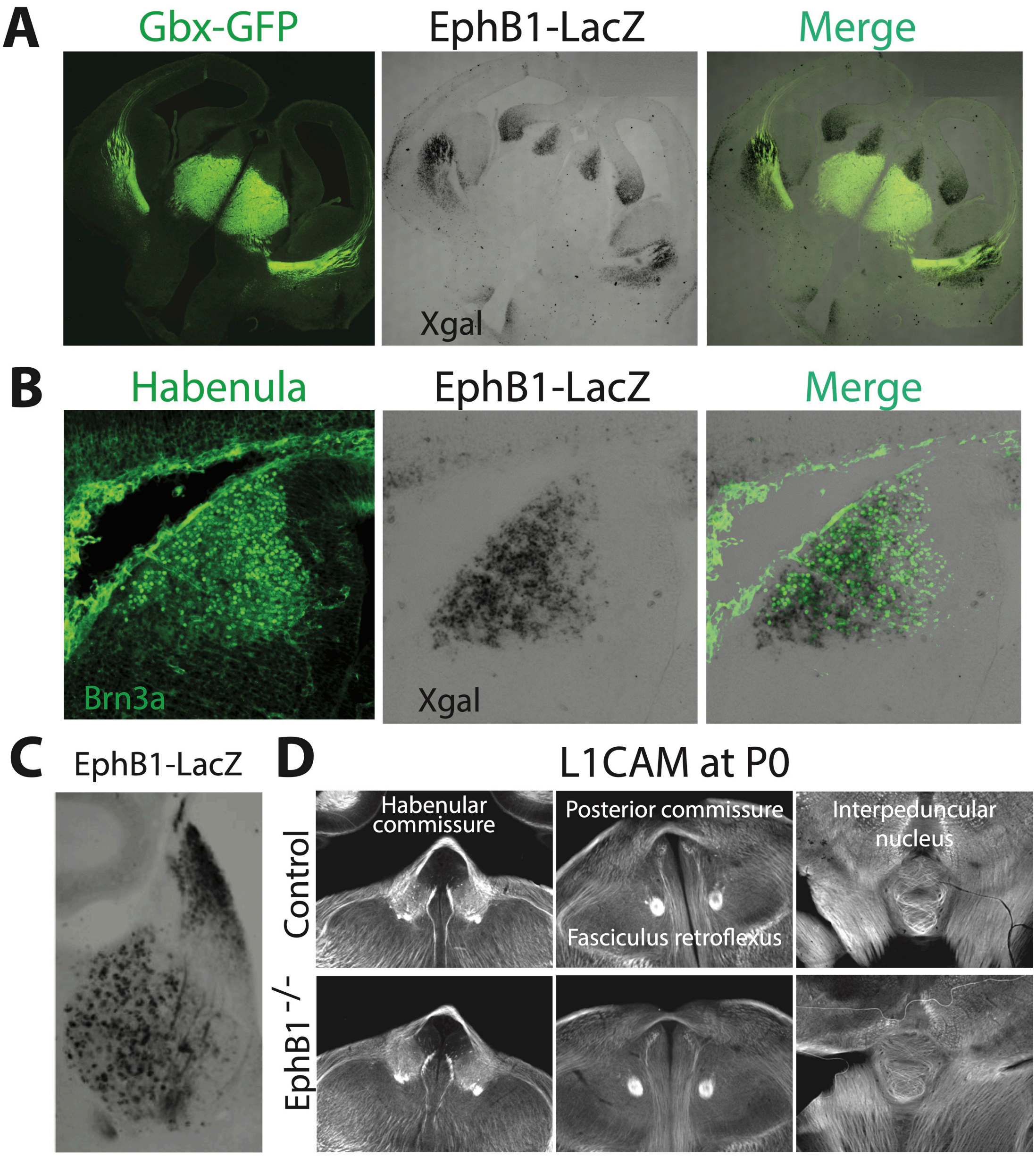


**Supplementary Figure 3**: **EphB1 expression in the habenula. A**. GFP and Xgal co-staining on coronal sections at E14.5 in Gbx-GFP;EphB1-LacZ embryos. **B**. Brn3a (marker of developing habenula) and Xgal co-staining on coronal sections of the habenula at E14.5 in EphB1-LacZ embryos. **C**. Xgal staining on coronal sections of the habenula in adult EphB1-LacZ mice. **D**. L1CAM staining on coronal sections of habenular axon tracts at P0 in control (upper panel in **D**) and EphB1$^{-/-}$ mice (lower panel in **D**). The images were taken using a microscope 10X objective.

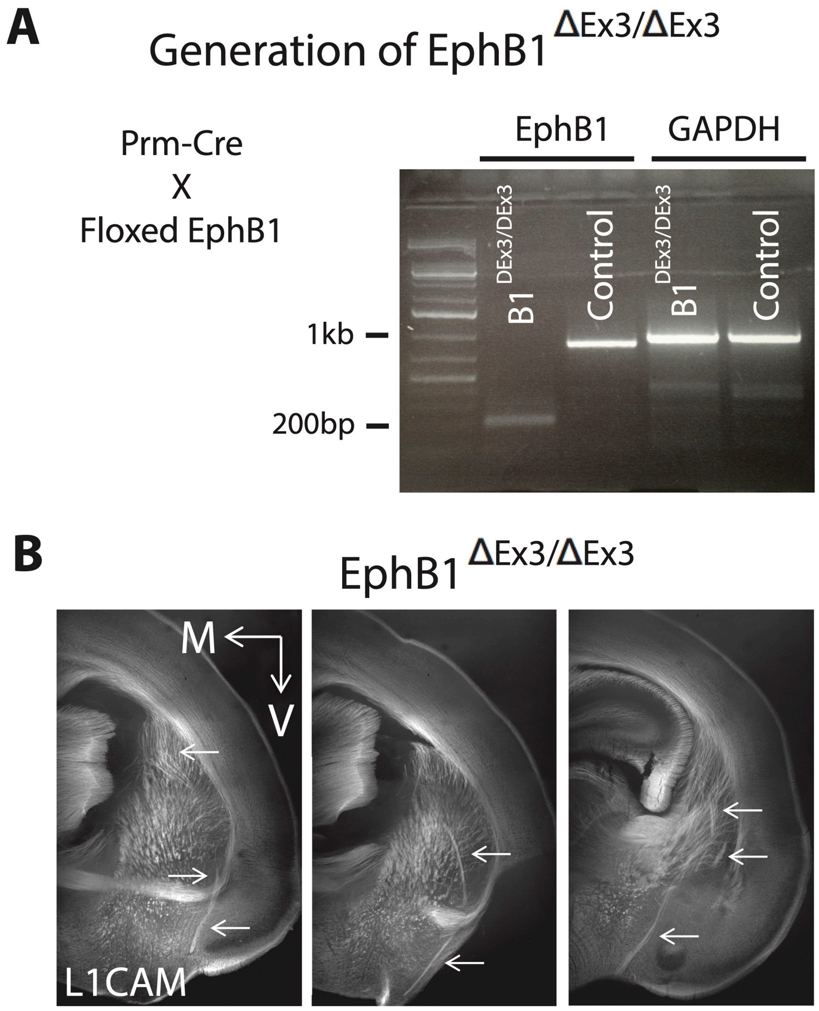


**Supplementary Figure 4**: **Generation and validation of floxed EphB1 mice**. **A**. Generation of a novel global EphB1 knockout mouse. EphB1$^{lox/lox}$ mice were crossed to Prm-Cre mice to generate germline transmission of the EphB1 loss-of-function allele (EphB1$^{deltaEx3/deltaEx3}$). RT-PCR showing efficient recombination of EphB1 exon 3 inEphB1$^{deltaEx3/deltaEx3}$ mice compared to control mice (excised exon 3 band: 153bp; control band: 880bp). GAPDH was used as a control. **B**. L1CAM staining on coronal sections at three different rostro-caudal levels at P0 on EphB1$^{deltaEx3/deltaEx3}$ mice, showing the same axon guidance defects (arrows) as in global EphB1$^{-/-}$ mice. The images were taken using a microscope 10X objective. V: ventral; M: medial.

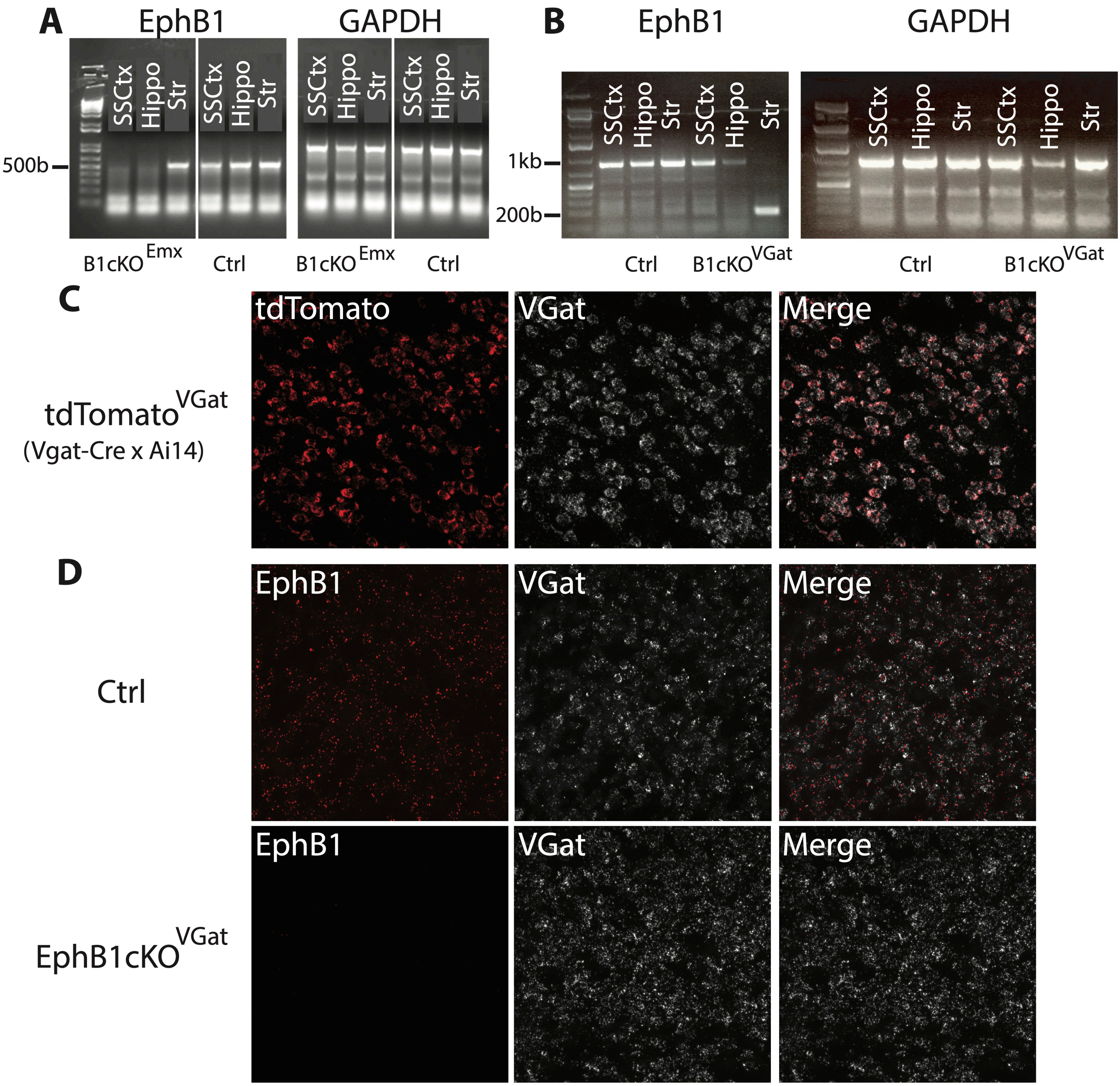


**Supplementary Figure 5**: **EphB1 deletion validation in EphB1 $cKO^{Emx}$ and in EphB1 $cKO^{Vgat}$ mice.**
**A, B**. RT-PCR showing specific recombination of EphB1 exon 3 in EphB1 $cKO^{Emx}$ mice (excised exon 3: no band; control band: 545bp) (**A**) and in EphB1 $cKO^{Vgat}$ mice (excised exon 3 band: 153bp; control band: 880bp) (**B**) compared to control mice. The two sets of primers for detection are described in the materials and methods section. GAPDH was used as a control. SSCtx: somatosensory cortex; Hippo: hippocampus; Str: striatum. **C**. tdTomato and Vgat co-staining using fluorescent in situ hybridization (RNAscope) on coronal sections of the dorsal striatum in $tdTomato^{Vgat}$ mice, showing perfect colocalization between tdTomato and endogenous Vgat. **D**. EphB1 and Vgat co-staining using RNAscope on coronal sections of the dorsal striatum in $tdTomato^{Vgat}$ control mice (upper panel in **D**) and in $tdTomato^{Vgat}$;EphB1 $cKO^{Vgat}$ mice (lower panel in $^{D}$), showing loss of EphB1 expression in Vgat-positive cells after EphB1 deletion. The images were taken using a confocal 20X objective.

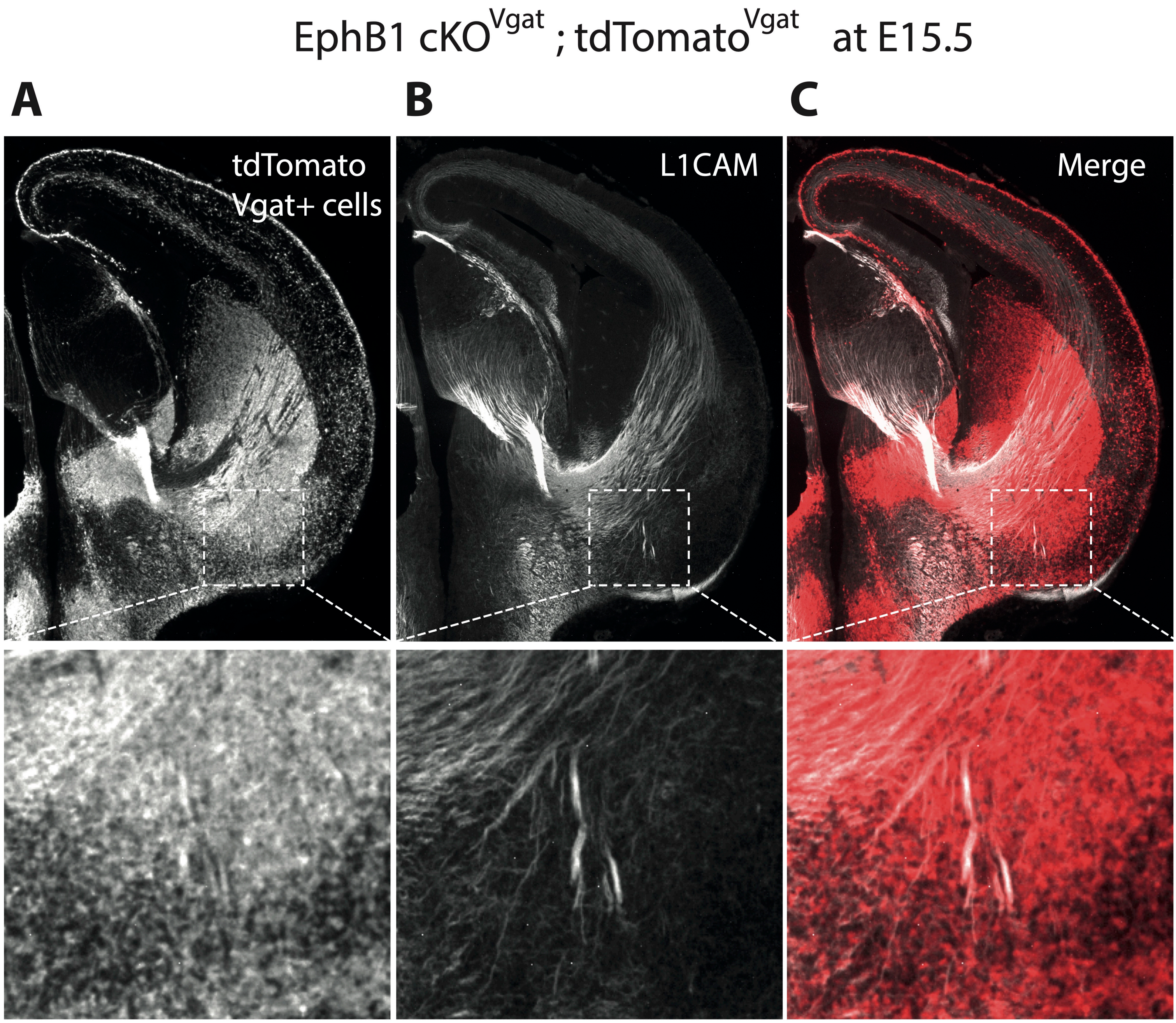


**Supplementary Figure 6**: **Vgat positive misguided axons at E15.5.** Ds-Red (**A**) and L1CAM (**B**) co-staining on coronal sections in EphB1 cKO$^{Vgat}$;tdTomato$^{Vgat}$ mice (Vgat-Cre x EphB1$^{lox/lox}$ x Ai14) at E15.5, showing Vgat positive misguided axons at E15.5 among L1CAM positive axons. The images in the lower panel are a zoom of the images in the white squares of the upper panel. The images were taken using a microscope 10X objective.

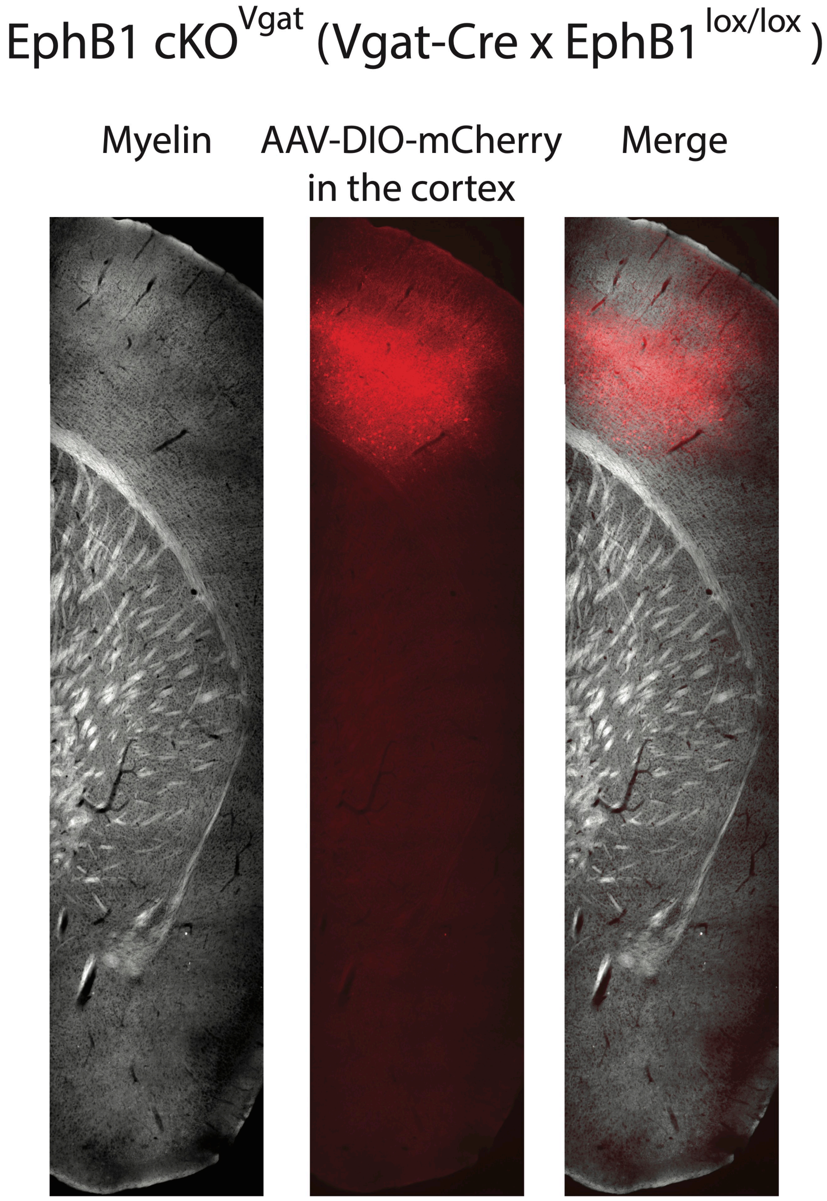


**Supplementary Figure 7**: **No misguided cortical long-range GABAergic projections in EphB1 cKOVgat mice.** Myelin and Ds-Red co-staining on coronal sections of adult EphB1 cKO$^{Vgat}$ mice, following Cre-dependent (DIO) mCherry AAV virus injections in the somatosensory cortex. The images were taken using a microscope 10X objective.

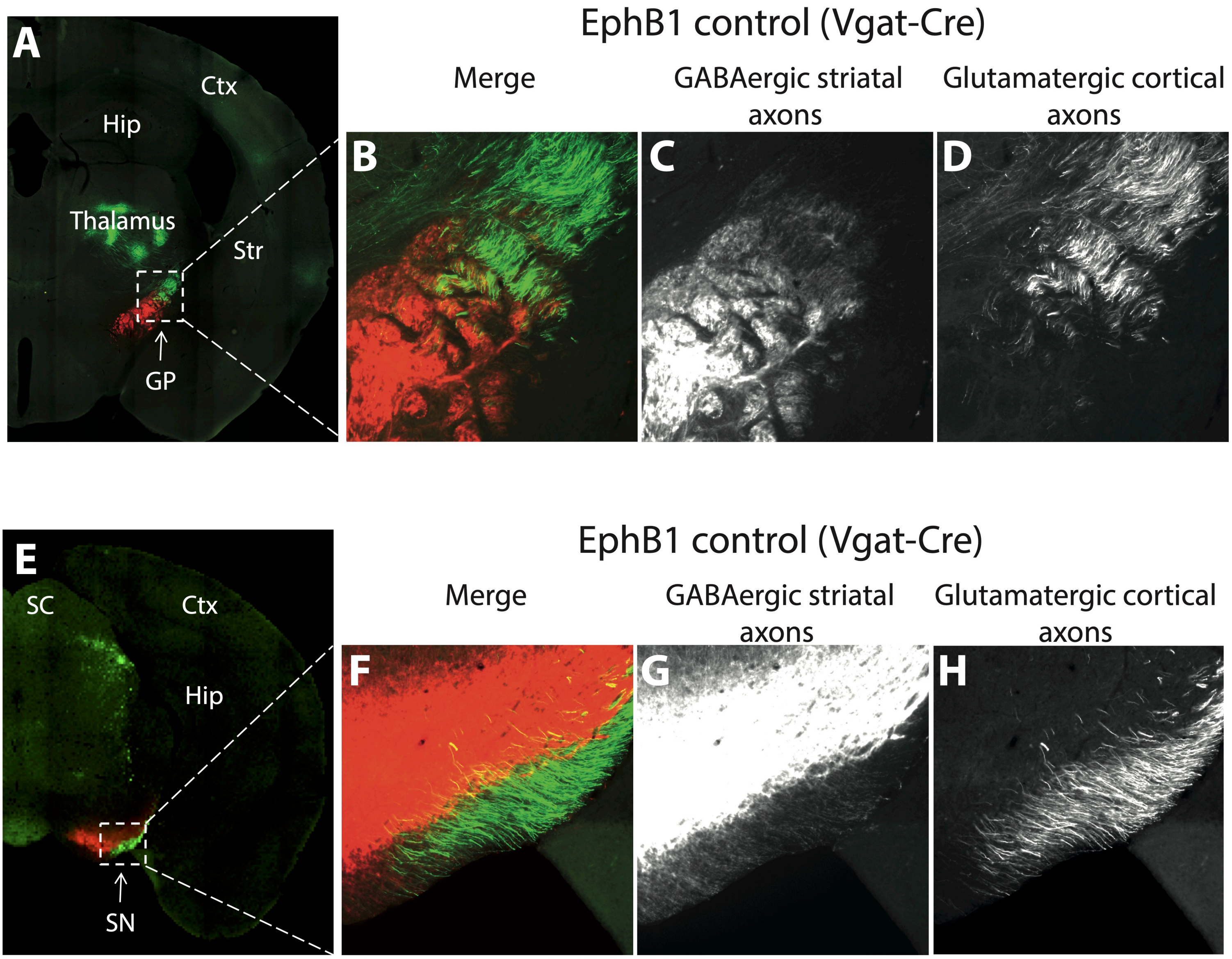


**Supplementary Figure 8: Cofasciculation of striatal GABAergic and somato-sensory-cortical glutamatergic axons**. Ds-Red (**C, G**) and GFP (**D, H**) co-staining on coronal sections in control mice (Vgat-Cre mice), at the level of the globus pallidus (**A-D**) and of the substantia nigra (**E-H**), following Cre-dependent (DIO) mCherry AAV virus injections in the dorsal striatum and CaMKII GFP AAV virus injections in the somatosensory cortex. The images were taken using a microscope 10X objective. Ctx: cortex; Hip: hippocampus; Str: striatum; GP: globus pallidus; SC: superior colliculus; SN: substantia nigra.

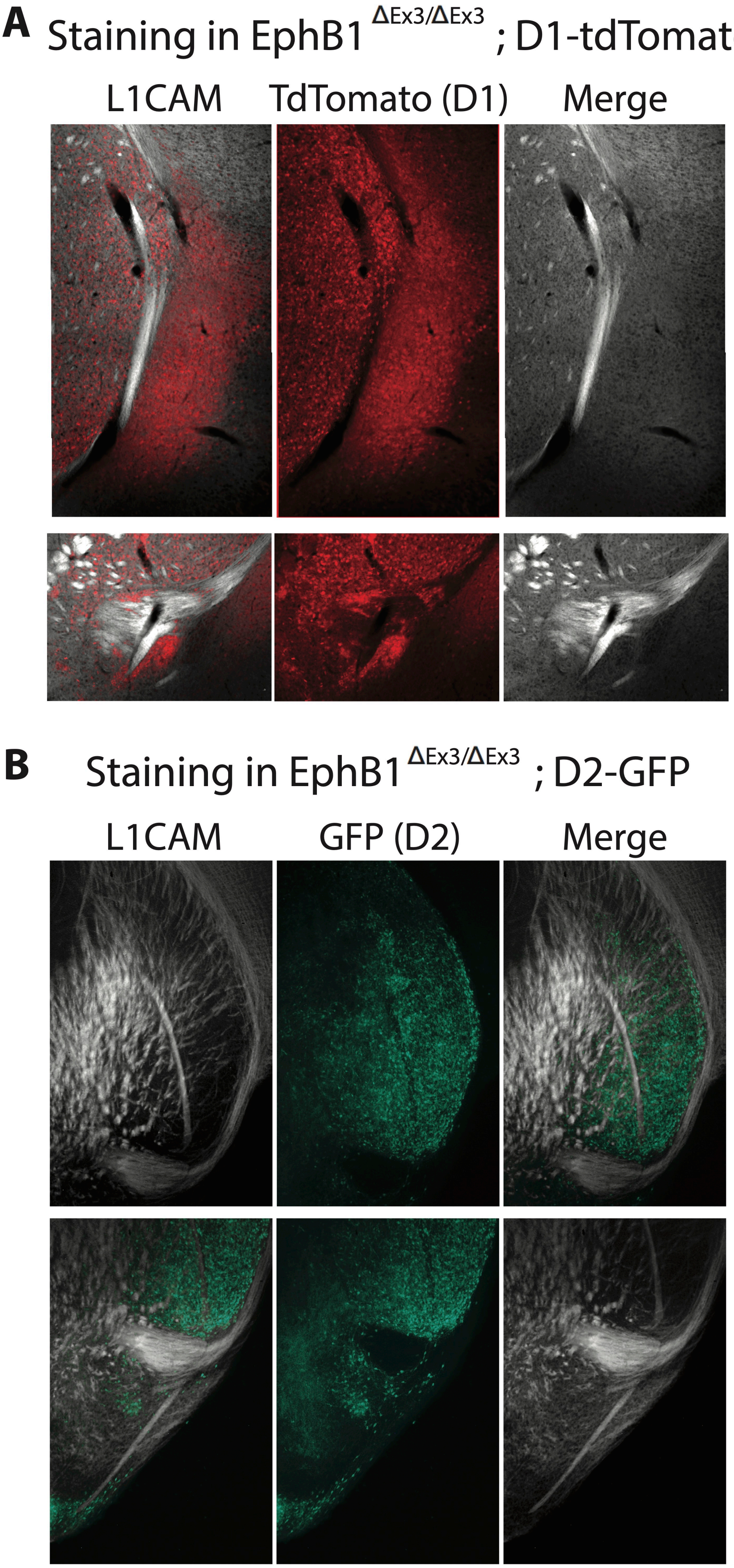


**Supplementary Figure 9**: **No clear misguided axons from D1- and D2-SPNs in EphB1$^{deltaEx3/deltaEx3}$ mice**. **A**. Myelin and Ds-Red co-staining on coronal sections of adult D1-tdTomato;EphB1$^{deltaEx3/deltaEx3}$ mice. **B**. L1CAM and GFP co-staining on coronal sections of D2-GFP;EphB1$^{deltaEx3/deltaEx3}$ pups at P0. The images were taken using a microscope 10X objective. V: ventral; M: medial.

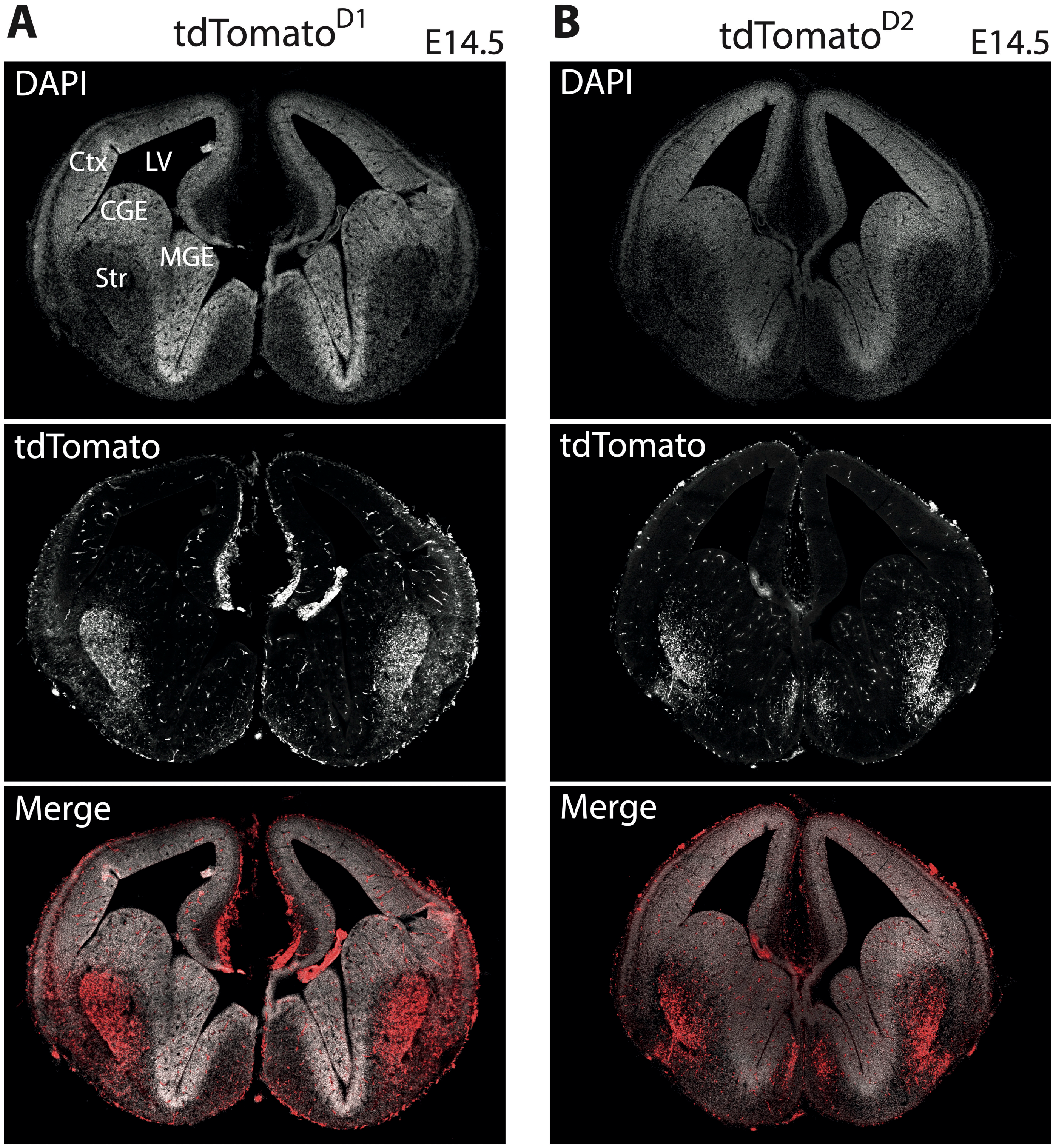


**Supplementary Figure 10: Effective Cre recombination at E14.5 of Drd1-Cre and Drd2-Cre mice.** DAPI and RFP (for tdTomato) co-staining on coronal sections in tdTomato$^{D1}$ (Drd1-Cre x Ai14; **A**) and tdTomato$^{D2}$ (Drd2-Cre x Ai14; **B**) mice at E14.5, showing effective recombination in the developing brain, with strong tdTomato expression in the developing striatum. The images were taken using a microscope 10X objective. LV: lateral ventricle; Ctx: cortex; CGE: caudal ganglionic eminence; MGE: medial ganglionic eminence; Str: striatum.

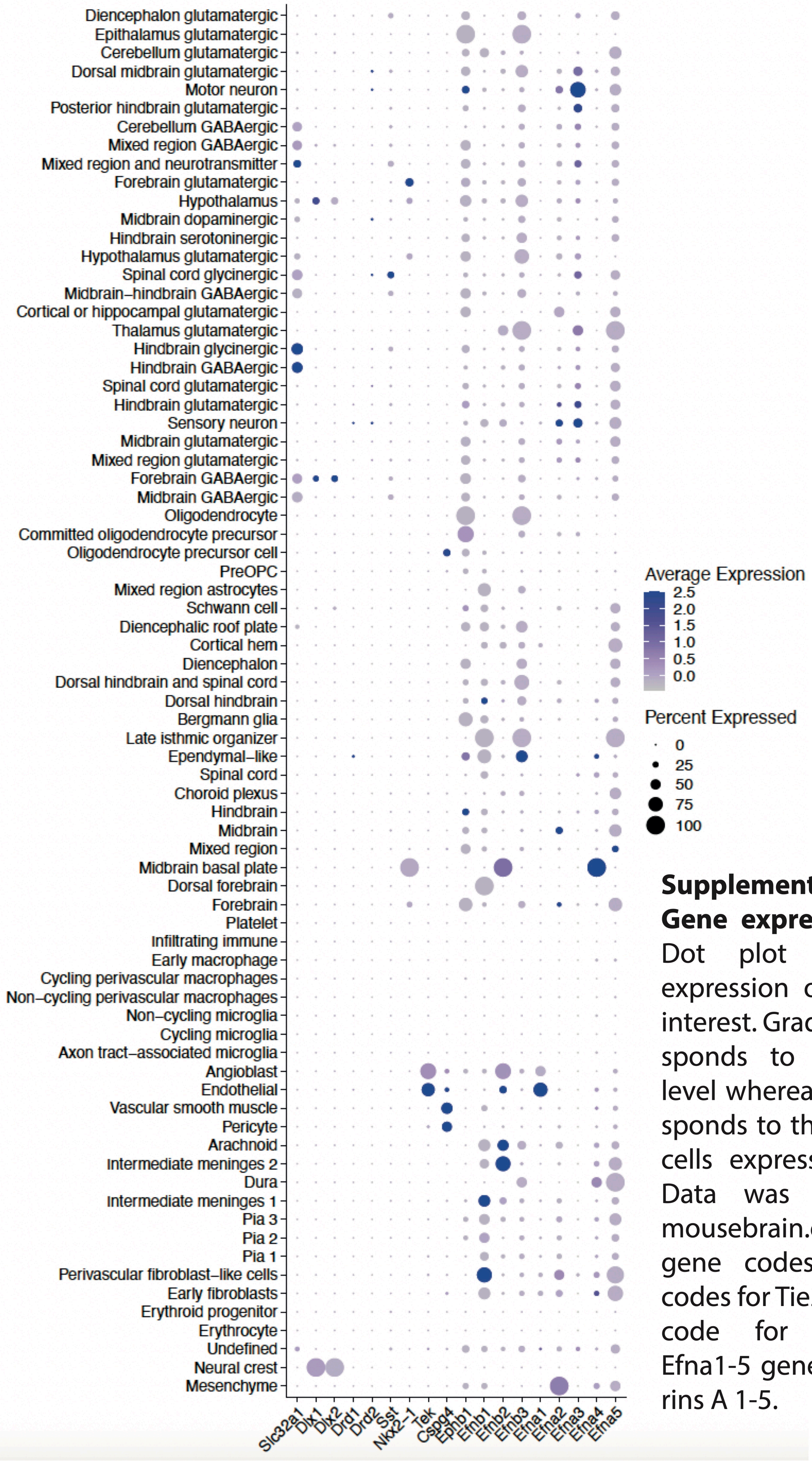


**Supplementary Figure 11: Gene expression at E14.5.** Dot plot depicting the expression of our genes of interest. Gradient color corresponds to the expression level whereas dot size corresponds to the percentage of cells expressing the genes. Data was retrieved from mousebrain.org. Slc32a1 gene codes for Vgat; Tek codes for Tie2; Efnb1-3 genes code for ephrins B1-3; Efna1-5 genes code for ephrins A 1-5.

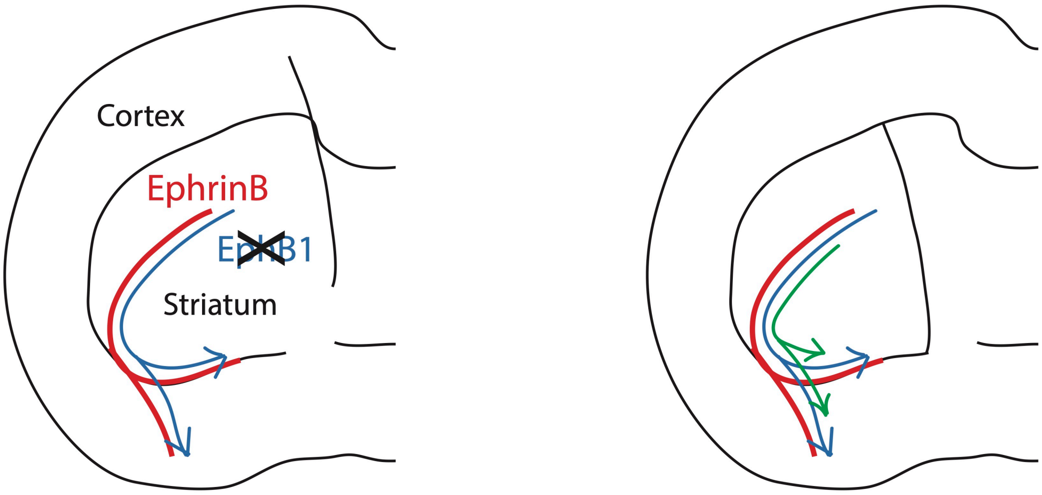


In absence of EphB1

1. Navigating GABAergic axons ectopically follow developing blood vessels

2. Misprojected GABAergic axons cause cortical axon misrouting through a cell non-autonomous mechanism

**Supplementary Figure 12: Hypothetical recapitulative scheme.**
